# The galectin-3 carbohydrate recognition domain maintains plasma membrane phosphatidylserine required for KRAS function

**DOI:** 10.64898/2026.10.30.763154

**Authors:** Kylie Rice, Byung-Seok Kim, Junchen Liu, John F. Hancock, Kwang-jin Cho

## Abstract

KRAS is a small GTPase essential for cellular proliferation, differentiation, and survival. It is predominantly localized to the plasma membrane (PM), where it forms nanoclusters essential for signal transduction. This PM binding and nanoclustering strictly depend on phosphatidylserine (PtdSer), an anionic phospholipid enriched in the inner PM leaflet. Galectin-3 (Gal-3), a β-galactoside-binding lectin implicated in cell growth, migration and tumor progression, has been shown to regulate KRAS PM binding and nanoclustering. Here, we demonstrate that *Gal-3* knockout (KO) or inhibition of its carbohydrate recognition domain (CRD) depletes KRAS and PtdSer from the PM. Exogenous PtdSer supplementation fully restores PM PtdSer and partially restores KRAS PM binding, indicating that Gal-3 promotes KRAS PM binding in part by maintaining PM PtdSer content. Total cellular PtdSer, PM PI4P, cholesterol and GM3 are unchanged, excluding these established regulators of PM PtdSer. Instead, *Gal-3* CRD inhibition disrupts the cellular balance of sphingomyelin (SM) and ceramide, alters intracellular SM distribution, and impairs the endocytic and recycling transport of PtdSer, SM, and epidermal growth factor receptor (EGFR). Together, our findings demonstrate that Gal-3 CRD activity sustains PM PtdSer content through endocytic trafficking, providing the PM lipid composition required for KRAS PM spatial organization.

## Introduction

K-, H- and NRAS proteins are small GTPases that cycle between an inactive GDP-bound and an active GTP-bound state, serving as key molecular switches in signaling pathways that control cell proliferation, survival and differentiation (Gorfe and Cho, 2021; Henkels et al., 2021). They are ubiquitously expressed in mammalian cells, and consistent with their central role in these processes, activating point mutations in RAS are found in approximately 18 – 20 % of human cancers, with *KRAS* mutations being the most prevalent (Prior et al., 2020). RAS proteins localize predominantly to the inner leaflet of the plasma membrane (PM), where they are spatially segregated into nanoscale clusters, called nanoclusters, essential for promoting high-fidelity signal transduction (Harding and Hancock, 2008; Tian et al., 2007). Stable PM association of H- and NRAS requires a series of post-translational modifications that generate lipid membrane anchors consisting of a C-terminal farnesyl moiety together with two additional palmitoyl moieties for HRAS and one for NRAS (Hancock et al., 1989; Hancock et al., 1990). For KRAS4B, the major splicing variant of KRAS (Hood et al., 2023) (hereafter KRAS), PM association relies on a C-terminal farnesyl moiety coupled with an adjacent polybasic domain composed of six consecutive lysine residues (Gutierrez et al., 1989; Hancock et al., 1991; Hancock et al., 1989; Hancock et al., 1990). This combination gives KRAS a unique binding specificity for phosphatidylserine (PtdSer) with one saturated and one unsaturated acyl chain (Zhou et al., 2017). This lipid-binding specificity is intrinsically encoded in the structure of the membrane anchor, making KRAS PM binding, nanoclustering, and thereby its biological function highly sensitive to the PM PtdSer content (Cho et al., 2012; Cho et al., 2013; Garrido et al., 2020; Lange et al., 2025; van der Hoeven et al., 2013; van der Hoeven et al., 2018).

PtdSer is synthesized in the endoplasmic reticulum (ER) and transported to the PM via lipid transfer proteins, oxysterol-binding protein-related protein (ORP) 5 and 8 at ER-PM membrane contact sites (Chung et al., 2015; Moser von Filseck et al., 2015). ORP5 and 8 form a heterodimer and exchange ER-derived PtdSer for PM phosphatidylinositol 4-phosphate (PI4P), which is synthesized by PI 4-kinase IIIα (PI4KA). Once transferred to the ER, PI4P is converted back to phosphatidylinositol (PI) by the phosphatase SAC1. Together, PI4KA and SAC1 maintain the PI4P gradient that enables the continuous delivery of PtdSer to the PM (Chung et al., 2015; Moser von Filseck et al., 2015). Genetic or chemical disruption of any part of this PtdSer/PI4P exchange machinery leads to a reduction in PM PtdSer levels, resulting in KRAS PM dissociation and impaired KRAS signaling (Gulyas et al., 2017; Kattan et al., 2021). Similarly, other strategies that reduce PM PtdSer content also dissociate KRAS from the PM and impair KRAS activity (Garrido et al., 2020; Lange et al., 2025; Miller et al., 2019; Rehl et al., 2023; Tan et al., 2019; Tan et al., 2018). In addition to this non-vesicular route, PtdSer cycles between the PM and endomembranes through endocytic and recycling transport, and intracellular PtdSer is itself required for retrograde traffic through endosomes (Kay and Fairn, 2019; Uchida et al., 2011). It is unclear how this vesicular transport is regulated, and the extent to which it sets steady-state PM PtdSer content.

Galectins are a family of carbohydrate-binding proteins defined by a β-galactoside-binding site within a carbohydrate recognition domain (CRD), typically located in the C-terminal region (Johannes et al., 2018). Galectins are involved in diverse biological processes including inflammation, immune responses, cell migration, autophagy and cell signaling (Johannes et al., 2018). They are localized primarily in the cytosol and nucleus, and are also secreted to the extracellular space via a non-classical pathway (Cummings et al., 2015). In addition to their canonical carbohydrate-binding function, galectins can interact with proteins lacking β-galactosides through non-carbohydrate-binding sites (Kim et al., 2013; Li et al., 2013). Previous studies reported that genetic loss of *galectin-3* (*Gal-3*), one of 15 galectin family members, blocks the PM binding, nanoclustering and signal output of KRAS, but not other RAS isoforms, whereas Gal-3 overexpression elevates KRAS nanoclustering and signal output (Belanis et al., 2008; Elad-Sfadia et al., 2004; Shalom-Feuerstein et al., 2008). While the exact mechanism of Gal-3-mediated KRAS PM binding has not been fully elucidated, studies have proposed that Gal-3 interacts with the farnesyl moiety of GTP-bound KRAS through its C-terminal CRD, and blocking this interaction dissociates KRAS from the PM (Elad-Sfadia et al., 2004; Levy et al., 2010; Shalom-Feuerstein et al., 2008). Separately, Gal-3 has an established role in endosomal membrane organization. Secreted Gal-3 re-enters cells by clathrin-independent endocytosis and cycles through endosomal and recycling compartments (Straube et al., 2013), where CRD-dependent oligomerization assembles a galectin lattice that drives the biogenesis of clathrin-independent carriers and the endosomal sorting of glycosylated cargo (Lakshminarayan et al., 2014; Nabi et al., 2015). Consistent with this, silencing Gal-3 delays the internalization and endosomal trafficking of EGFR (Liu et al., 2012).

In this study, we have discovered that Gal-3-regulated KRAS PM localization and nanoclustering are mediated by maintaining PM PtdSer content. We demonstrate that *Gal-3* knockout (KO) or inhibiting the CRD depletes KRAS and PtdSer from the PM, and that exogenous PtdSer supplementation fully restores PM PtdSer and partially restores KRAS. *Gal-3* KO and CRD inhibition also perturb the metabolism of sphingomyelin (SM) and ceramide (Cer), and alter the intracellular distribution of SM. We further show that *Gal-3* KO delays the endocytic and recycling transport of PtdSer, SM and epidermal growth factor receptor (EGFR). We propose that Gal-3 CRD activity maintains PM PtdSer content by supporting the endocytic trafficking of PtdSer and SM, and that the resulting loss of PM PtdSer disrupts KRAS PM binding and nanoclustering.

## RESULTS

### *Gal-3* knockout depletes KRAS and PtdSer from the PM

To dissect the mechanism by which Gal-3 promotes KRAS PM binding and nanoclustering (Shalom-Feuerstein et al., 2008), we examined whether Gal-3 regulates the PM localization of PtdSer. *Gal-3* was knocked out using CRISPR/Cas9 in Madin-Darby canine kidney (MDCK) cells stably expressing GFP-tagged oncogenic KRAS (KRASG12V) or -LactC2, a well-characterized PtdSer marker (Cho et al., 2012; Yeung et al., 2008). We used two Gal-3-targeting single guide RNA (sgRNA), sets 2 and 3, which introduced nucleotide insertions, resulting in premature termination codons at amino acid residues 242 and 200, respectively, within the CRD of Gal-3 (Figs. 1A and B) (Agrwal et al., 1993; Birdsall et al., 2001; Miller et al., 2016; Seetharaman et al., 1998). Immunoblotting using an anti-Gal-3 antibody targeting the N-terminal 18 amino acids showed undetectable Gal-3 protein levels, confirming *Gal-3* KO (Fig. 1C). To examine the PM localization of KRAS and PtdSer, we repeated *Gal-3* KO using these sgRNAs in MDCK cells stably co-expressing mCherry-CAAX, a generic endomembrane marker (Cho et al., 2012; Choy et al., 1999), together with GFP-KRASG12V or -LactC2. Our confocal microscopy showed that GFP-KRASG12V was redistributed from the PM (Fig. 1D). Manders’ coefficient values, which measure the fraction of mCherry-CAAX co-localized with GFP-KRASG12V, were increased after *Gal-3* KO, suggesting KRAS redistribution to endomembranes. GFP-LactC2 showed a similar trend, redistributed to endomembranes (Fig. 1E). Our electron microscopy (EM) analysis of intact basal PM sheets prepared from these cells and labeled with anti-GFP antibodies coupled to 4.5-nm gold particles showed that the number of gold particles, which correlates with the concentration of KRASG12V or with PtdSer content in the inner PM leaflet, was reduced, indicative of KRAS and PtdSer PM dissociation after *Gal-3* KO (Fig. 1F). Spatial mapping analysis further showed a reduction in the nanoclustering of both KRASG12V and PtdSer remaining at the PM (Fig. 1G), consistent with the previous study (Shalom-Feuerstein et al., 2008). Moreover, an annexin V-binding assay analyzed by flow cytometry showed that PM PtdSer was not externalized to the outer leaflet after *Gal-3* KO, excluding PtdSer scrambling or apoptosis as the cause of the reduced GFP-LactC2 PM localization (Fig. 1H). Together, our data suggest that Gal-3 maintains the PM localization of KRAS and PtdSer.

**Figure 1.**
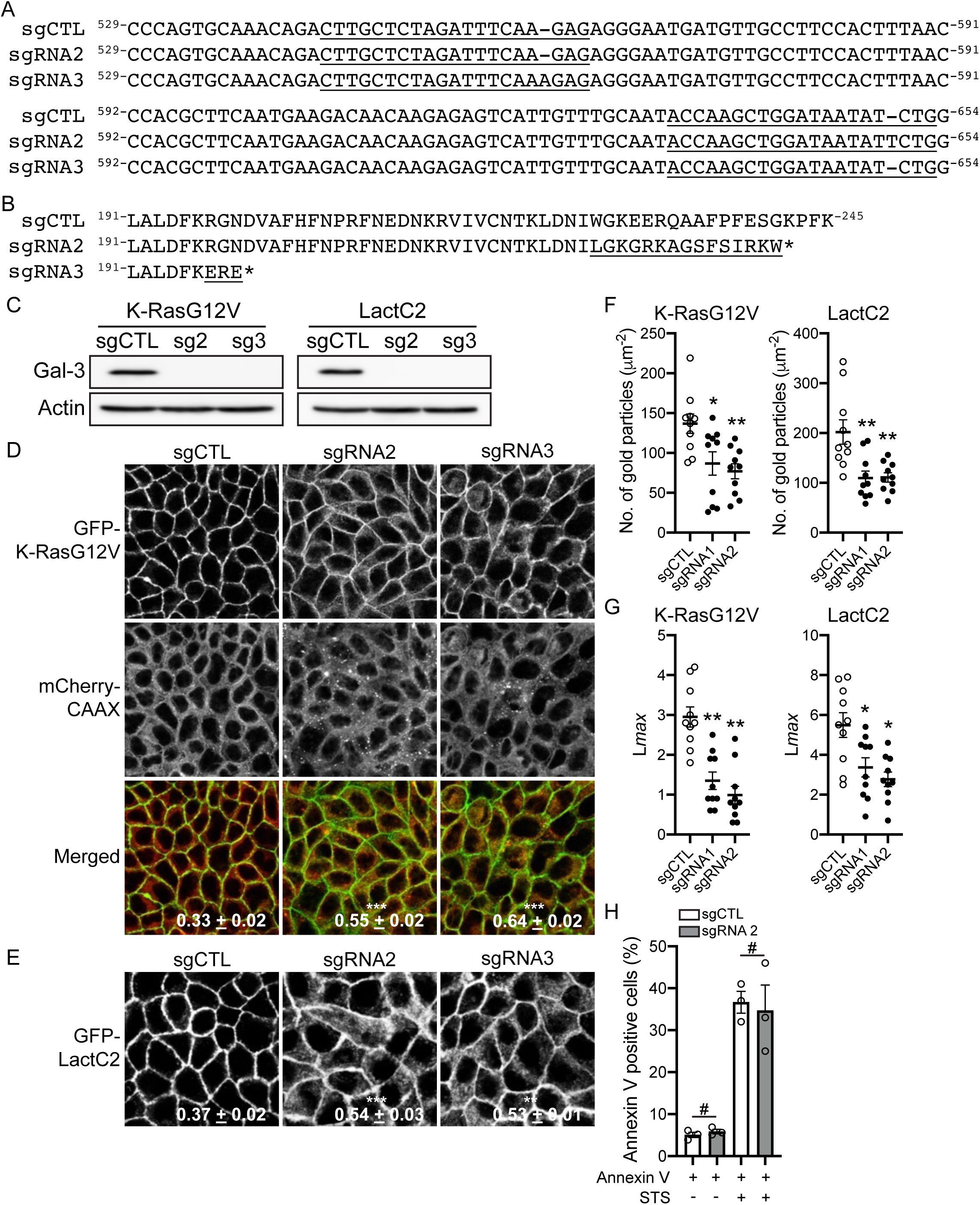
*Gal-3* KO depletes KRAS and PtdSer from the PM. **(A)** Alignment of the endogenous *Canis lupus familiaris* Gal-3 nucleotide sequence (NM_001197043.1). Underlined nucleotides indicate the Gal-3 sequence targeted by a Gal-3-specific sgRNA. A thymine (sgRNA 2) or an adenine (sgRNA 3) was inserted at nucleotide position 651 or 561, respectively. An empty CRISPR/Cas9 vector was used as a control (sgCTL). **(B)** Alignment of endogenous canine Gal-3 amino acid sequence. Underlined amino acids indicate the frameshifted sequence downstream of each insertion, which introduces a premature termination codon within the Gal-3 CRD. **(C)** *Gal-3* was knocked out in MDCK cells stably co-expressing mCherry-CAAX and GFP-KRASG12V or - LactC2 and cell lysates were blotted for Gal-3. MDCK cells expressing non-targeting sgRNA were used as a control (sgCTL). Actin blots are shown as loading controls. **(D and E)** *Gal-3* KO MDCK cells co-expressing mCherry-CAAX and GFP-KRASG12V or - LactC2 were seeded on glass coverslips, fixed with 4% PFA and imaged by confocal microscopy. Values represent mean ± S.E.M. (n = 3) of the fraction of mCherry fluorophore co-localized with GFP fluorophore, calculated by Manders’ coefficient. mCherry-CAAX and merged images for (E) are shown in Fig. S1. Representative images from three independent experiments are shown. Scale bars = 10 μm. **(F)** Intact basal PM sheets prepared from these cells were labeled with 4.5 nm gold-conjugated anti-GFP antibodies and visualized by EM. The graph shows a mean number of gold particles ± S.E.M. (n ≥ 15). Representative images are shown in Fig. S1. **(G)** Spatial mapping of the same gold-labeled PM sheets was performed. The peak values, Lmax, of the respective weighted mean K-function *L(r) — r* curves are shown as bar graphs (n ≥ 15). **(H)** *Gal-3* KO MDCK cells stably co-expressing mCherry-CAAX and GFP-KRASG12V were incubated with Cy5-conjugated annexin V, and annexin V-positive cells were counted in a cytometer. Cells treated with 1 μM staurosporine for 6 h were used as positive controls to induce apoptosis. Statistical differences between control (sgCTL) and *Gal-3* KO cells were assessed using one-way ANOVA test for (D – F), bootstrap tests for (G) and Student’s t-test for (H) (* p < 0.05, ** p < 0.01, *** p < 0.001, # - not significant).

### Gal-3 CRD is required for PM PtdSer localization

KRAS interacts with the Gal-3 CRD to promote KRAS PM binding and nanoclustering (Shalom-Feuerstein et al., 2008). To examine whether Gal-3 CRD is required for PM PtdSer localization, we treated MDCK cells stably co-expressing mCherry-CAAX and GFP-KRASG12V or - LactC2 with G3-A9 or G3-C12, small peptides that bind and block Gal-3 CRD (Yang et al., 2014; Zou et al., 2005). Our confocal microscopy data showed that GFP-KRASG12V and -LactC2 were redistributed to endomembranes, indicated by elevated Manders’ coefficient values (Fig. 2A). EM analyses further revealed that blocking Gal-3 CRD reduced the PM localization and nanoclustering of KRAS and PtdSer (Figs. 2B and C). Next, we conducted PtdSer addback experiments to determine whether *Gal-3* KO or CRD inhibition dissociates KRAS from the PM by depleting PM PtdSer content. Supplementation with exogenous PtdSer fully restored the PM localization of PtdSer, while it partially restored KRASG12V (Fig. 3A). Together with Fig. 1, our data suggest that Gal-3 regulates the PM localization of PtdSer via the CRD, and that this at least partially accounts for its promotion of KRAS PM localization.

**Figure 2.**
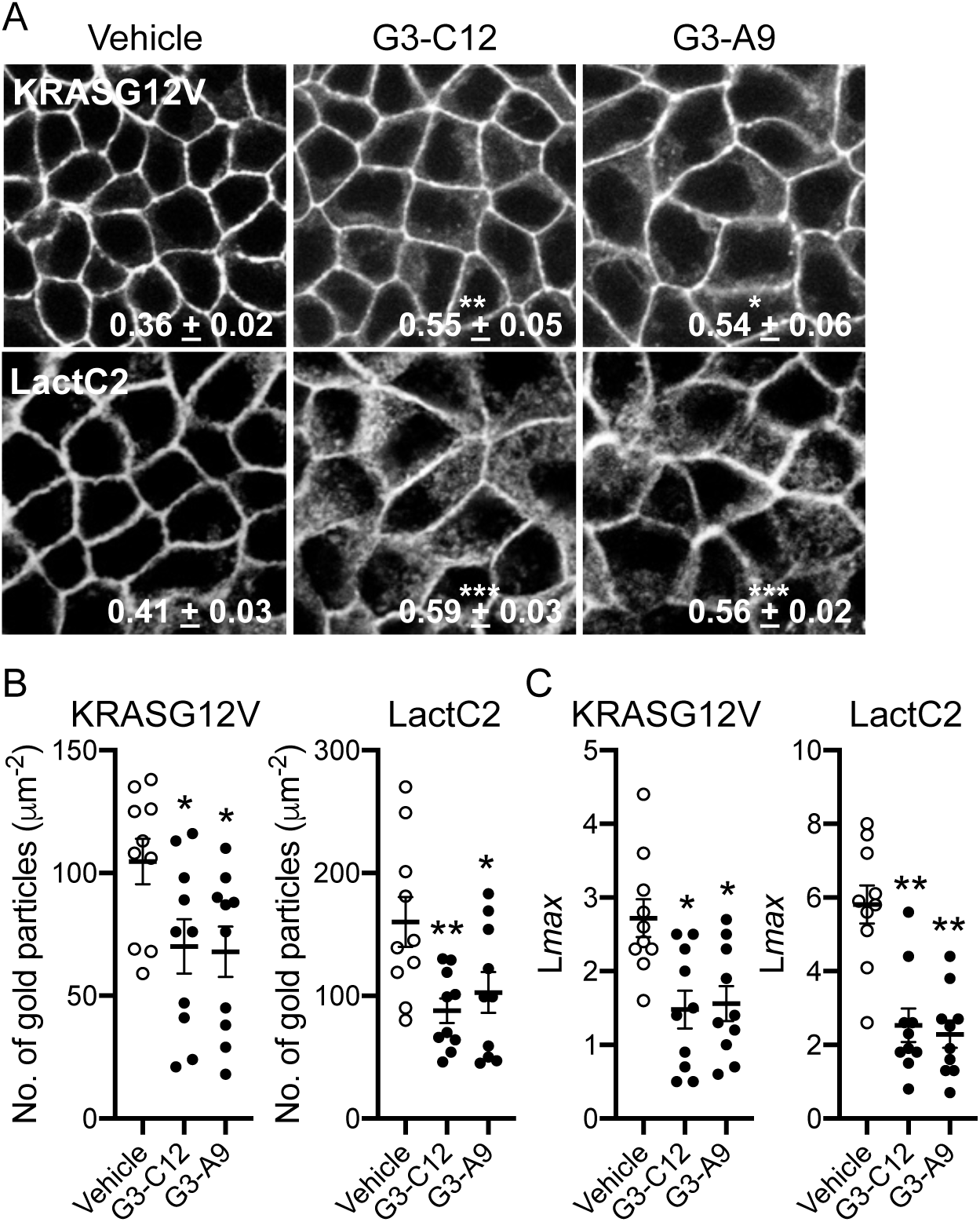
Gal-3 CRD inhibitors deplete KRAS and PtdSer from the PM. **(A)** MDCK cells stably co-expressing mCherry-CAAX and GFP-KRASG12V or -LactC2 were treated with 50 μM G3-A9 or G3-C12 for 48 h, fixed with 4% PFA and imaged by confocal microscopy. Representative images taken in the green channel only are shown. Values represent mean ± S.E.M. (n = 3) of the fraction of mCherry fluorophore co-localized with GFP fluorophore, calculated by Manders’ coefficient. Representative images from three independent experiments are shown. Scale bars = 10 μm. **(B)** Intact basal PM sheets prepared from these cells were labeled with 4.5 nm gold-conjugated anti-GFP antibodies and visualized by EM. The graph shows a mean number of gold particles ± S.E.M. (n ≥ 15). **(C)** Spatial mapping of the same gold-labeled PM sheets was performed. The peak values, Lmax, of the respective weighted mean K-function *L(r) — r* curves are shown as bar graphs (n ≥ 15). Statistical differences between vehicle-(control) and inhibitor-treated cells were assessed using one-way ANOVA tests for (A and B) and bootstrap tests for (C) (* p < 0.05, ** p < 0.01).

**Figure 3.**
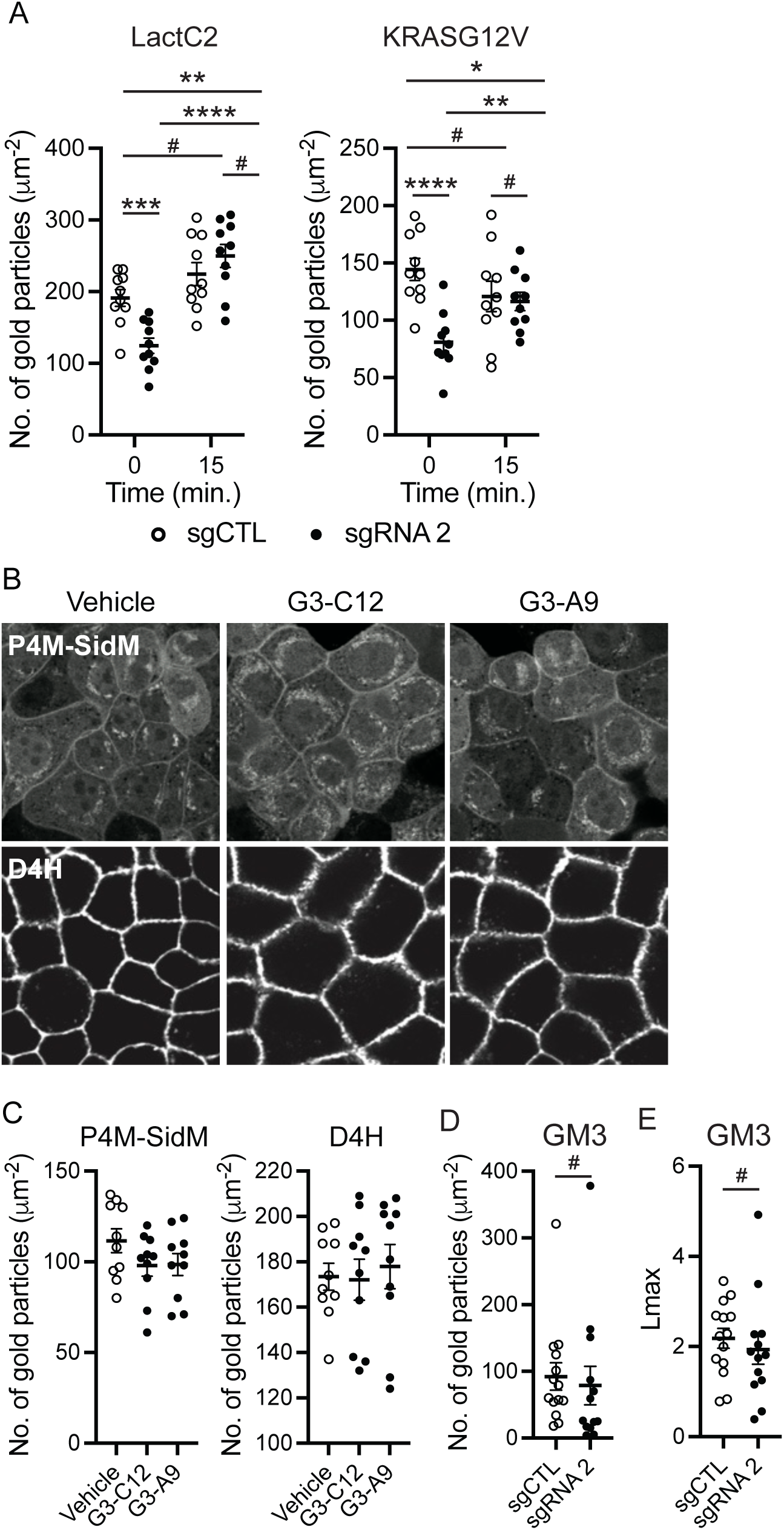
Exogenous PtdSer partially restores KRAS PM localization, and Gal-3 CRD inhibition does not alter PM PI4P, cholesterol or GM3. (A) *Gal-3* KO MDCK cells stably expressing GFP-LactC2 or -KRASG12V were incubated with 10 μM exogenous PtdSer for 15 min. Intact basal PM sheets were prepared from these cells, labeled with 4.5 nm gold-conjugated anti-GFP antibodies, and visualized by EM. Graphs show the mean number of gold particles ± S.E.M. (n ≥ 15). (B) MDCK cells stably expressing GFP-P4M-SidM or -D4H were treated with Gal-3 CRD inhibitors for 48 h, fixed with 4% PFA and imaged by confocal microscopy. Representative images from three independent experiments are shown. Scale bars = 10 μm. (C) Intact basal PM sheets prepared from these cells were labeled with 4.5 nm gold-conjugated anti-GFP antibodies and visualized by EM. The graph shows the mean number of gold particles ± S.E.M. (n ≥ 15). **(D)** Intact basal PM sheets from *Gal-3* KO MDCK cells stably expressing GFP-KRASG12V were prepared, labeled with 4.5 nm gold-conjugated anti-GM3 antibodies, and imaged by EM. The graph shows the mean number of gold particles ± S.E.M. (n ≥ 14). **(E)** Spatial mapping of the same gold-labeled PM sheets was performed. Peak values, Lmax, of the respective weighted mean K-function, *L(r) — r* curves, are shown as bar graphs (n ≥ 14). Statistical differences between control (sgCTL) and *Gal-3* KO cells, as well as before and after addback, were assessed by Student’s t-tests for (A and D) and bootstrap tests for (E). Statistical differences between vehicle- and Gal-3 CRD inhibitor-treated cells were assessed by one-way ANOVA tests for (C). (* p < 0.05, ** p < 0.01, *** p < 0.001, **** p < 0.0001, # - not significant).

### *Gal-3* KO or CRD inhibition does not alter PM PI4P, cholesterol or GM3 content

We have previously reported that a pool of PtdSer co-localizes with cholesterol in the PM, and that their PM localizations are interdependent (Cho et al., 2016). Also, PM PI4P content is essential for PM PtdSer localization via the lipid-transfer proteins ORP5 and 8 (Chung et al., 2015; Moser von Filseck et al., 2015). Moreover, our recent study has demonstrated that elevating PM PI3P content by blocking PI3P to PI conversion depletes PM PI4P content, thereby reducing PM PtdSer (Lange et al., 2025). To examine whether Gal-3 CRD maintains PM PtdSer content via these mechanisms, MDCK cells stably expressing the lipid probes GFP-P4M-SidM (Hammond et al., 2014) and GFP-D4H (Maekawa and Fairn, 2015), which report PI4P and cholesterol respectively, were treated with the Gal-3 CRD blockers, and their cellular localizations were analyzed by confocal microscopy and EM. Our data showed that the inhibitors did not change the PM localization of these probes (Figs. 3B and C), suggesting that PM PtdSer depletion upon Gal-3 CRD inhibition is independent of these PM lipids. Another set of lipids that regulate the PM localization of PtdSer and KRAS comprises sphingomyelin (SM) and ceramide (Cer). Perturbing the metabolism of SM or Cer blocks the PM localization of PtdSer and KRAS (Cho et al., 2016; Tan et al., 2018; van der Hoeven et al., 2018). Also, Cer is a building block for GM3 (monosialodihexosylganglioside) and SM4 (sulfatide), glycosphingolipids predominantly found in the outer PM leaflet (Ryckman et al., 2020). A recent study has reported that genetic or metabolic disruption of GM3 or SM4 biosynthesis impaired the PM localization of PtdSer and KRAS (Liu et al., 2023). The authors propose that GM3 and SM4 in the outer PM leaflet mediate cross-bilayer coupling with PtdSer in the inner PM leaflet to maintain PM PtdSer content, and that loss of these glycosphingolipids promotes PtdSer and KRAS dissociation from the PM (Liu et al., 2023). To examine whether Gal-3 regulates the PM localization of PtdSer and KRAS through reduced PM GM3 levels, we directly measured the PM content of GM3 in *Gal-3* KO cells by EM using anti-GM3 immunogold labeling (Liu et al., 2023). Our data showed that GM3 levels and nanoclustering in the outer PM leaflet were not changed (Figs. 3D and E). Together, our data suggest that *Gal-3* KO or CRD inhibition blocks the PM localization of PtdSer and KRAS via mechanisms that are independent of PM PI4P, cholesterol and GM3.

### *Gal-3* KO or CRD inhibition disrupts the metabolism of sphingomyelin and ceramide

To determine whether Gal-3 CRD inhibition reduces total PtdSer levels, we performed whole cell lipidomics in Gal-3 CRD inhibitor-treated MDCK cells. We also measured total SM and Cer levels, since perturbing their metabolism blocks PM PtdSer localization (Cho et al., 2016; Tan et al., 2018; van der Hoeven et al., 2018). Our data showed that while Gal-3 CRD inhibitor treatment did not change total PtdSer levels, SM and Cer levels were elevated and reduced, respectively (Figs. 4A and B). In addition to these lipids, *Gal-3* KO and CRD inhibition changed other cellular lipids (Fig. S2). These data suggest that Gal-3 CRD inhibition alters the levels of multiple cellular lipid classes.

**Figure 4.**
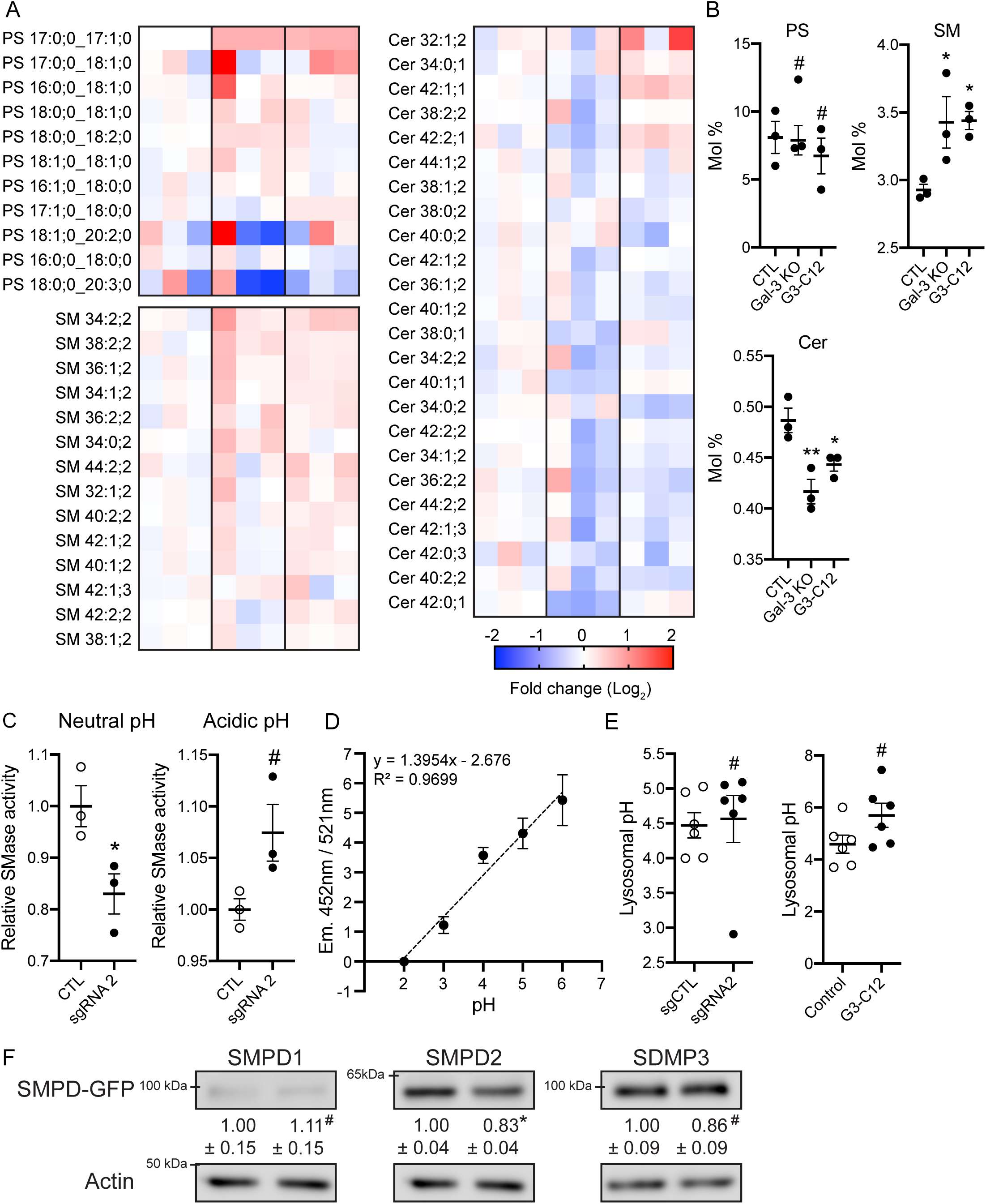
*Gal-3* KO and CRD inhibition disrupt the SM/Cer cellular balance. **(A)** MDCK cells stably expressing GFP-KRASG12V that are *Gal-3* KO or treated with a Gal-3 inhibitor for 48 h were prepared and whole cell SM, PtdSer and Cer levels were measured from three independent experiments. A heat map was constructed to quantify the changes of different lipid species in *Gal-3* KO or CRD inhibited cells. Each row represents a different species of lipid, while each column represents a single sample. The scaled expression values of each lipid measured are plotted in red-blue color log₂ scale. In relation to control cells, red and blue colored tiles indicate higher and lower lipid contents, respectively. **(B)** The graphs show the mean of total moles % ± S.E.M. (n = 3) of indicated lipids. Significant differences between control and Gal-3 CRD-inhibited cells were assessed using one-way ANOVA tests. **(C)** Cell lysates from *Gal-3* KO MDCK cells stably expressing GFP-KRASG12V were used to measure endogenous SMase activity. The assay was performed in a neutral (pH 7.5) or an acidic (pH 5.0) environment to measure specific activities of neutral SMase and acid SMase, respectively. The graph shows the mean conversion of SM to Cer ± S.E.M from three independent experiments. Significant differences between control (sgCTL) and *Gal-3* KO cells were assessed using Student’s t-tests (* p < 0.05; # - not significant). **(D)** WT MDCK cells were grown in the presence of Lysosensor Yellow/Blue dextran probe for 48 h and equilibrated with MES calibration buffer (pH 2 – 6) for 15 min at room temperature. Emissions of the probe at 452 nm and 521 nm were measured by a fluorescent plate reader using 355 nm and 381 nm excitation wavelengths, respectively. The mean 452/521 nm fluorescence ratio ± S.E.M. was plotted against pH from three independent experiments, and a best fit line was plotted. **(E)** *Gal-3* KO MDCK cells or WT MDCK cells treated with a CRD inhibitor for 48 h were grown in the presence of Lysosensor Yellow/Blue dextran probe. Emissions at 452 and 521 nm were measured by a fluorescent plate reader using 355 and 381 nm excitation wavelengths, respectively. The 452/521 nm fluorescence ratio ± S.E.M. from three independent experiments was used to determine the lysosomal pH using the calibration graph from (D). **(F)** MDCK cells stably expressing SMPD1-, SMPD2-, or SMPD3-GFP were treated with 50 μM G3-C12 for 48 h, and cell lysates were immunoblotted with an anti-GFP antibody to measure the total SMPD-GFP expression levels. Values indicate the relative mean ± S.E.M. of SMPD expression levels from three independent experiments. Actin blots are shown as loading controls. Significant differences between vehicle- (control) and G3-C12-treated cells were assessed using Student’s t-tests (* p < 0.05, # - not significant).

To examine whether Gal-3 regulates SM conversion to Cer, we measured the activities of acid or neutral sphingomyelinase (SMase) in *Gal-3* KO cells. Briefly, cell lysates of *Gal-3* KO MDCK cells were incubated with exogenous SM to measure the conversion of SM to Cer and phosphorylcholine by endogenous SMases. Phosphorylcholine is then hydrolyzed to choline, which in turn is oxidized, generating H₂O₂, a readout for SM to Cer conversion (Garrido et al., 2020; Tan et al., 2019). Furthermore, since SMase converts SM to Cer in acidic or neutral cellular pH environments by acid SMase (SMPD1) and neutral SMases (SMPD2 and SMPD3), respectively (Callahan et al., 1983; Liu et al., 1998; Marchesini and Hannun, 2004), we measured SMase activity in acidic (pH 5.0) and neutral (pH 7.5) environments. Our data showed that *Gal-3* KO reduced SMase activity at neutral but not acidic pH (Fig. 4C), while *Gal-3* KO did not alter lysosomal pH (Figs. 4D and E). Moreover, protein levels of GFP-SMPD2, but not of other SMases, were reduced in Gal-3 CRD-inhibited cells (Fig. 4F). These data suggest that Gal-3 CRD maintains SM/Cer balance by promoting neutral SMase activity.

### *Gal-3* KO and CRD inhibition perturb intracellular SM distribution

To further characterize the regulatory roles of Gal-3 in cellular SM, we analyzed its cellular distribution. *Gal-3* KO MDCK cells with or without permeabilization were stained with a GFP-tagged SM probe, the nontoxic recombinant fragment of lysenin (GFP-lysenin) (Cho et al., 2016; Maekawa et al., 2016; van der Hoeven et al., 2018). Our confocal microscopy showed that GFP-lysenin stained the outer leaflet of the PM, the major site of SM localization (Lange et al., 1989; Molitoris and Simon, 1985; Shiao and Vance, 1993; van Helvoort et al., 1997), in non-permeabilized control and *Gal-3* KO cells (Figs. 5A and B). In permeabilized control cells, GFP-lysenin was localized to intracellular vesicular structures, whereas it was more extensively distributed in elongated structures at the perinuclear region in *Gal-3* KO cells (Figs. 5C and D). A similar effect was observed with the CRD inhibitors (Figs. 5E and F). These data suggest that Gal-3 CRD regulates the intracellular distribution of SM.

**Figure 5.**
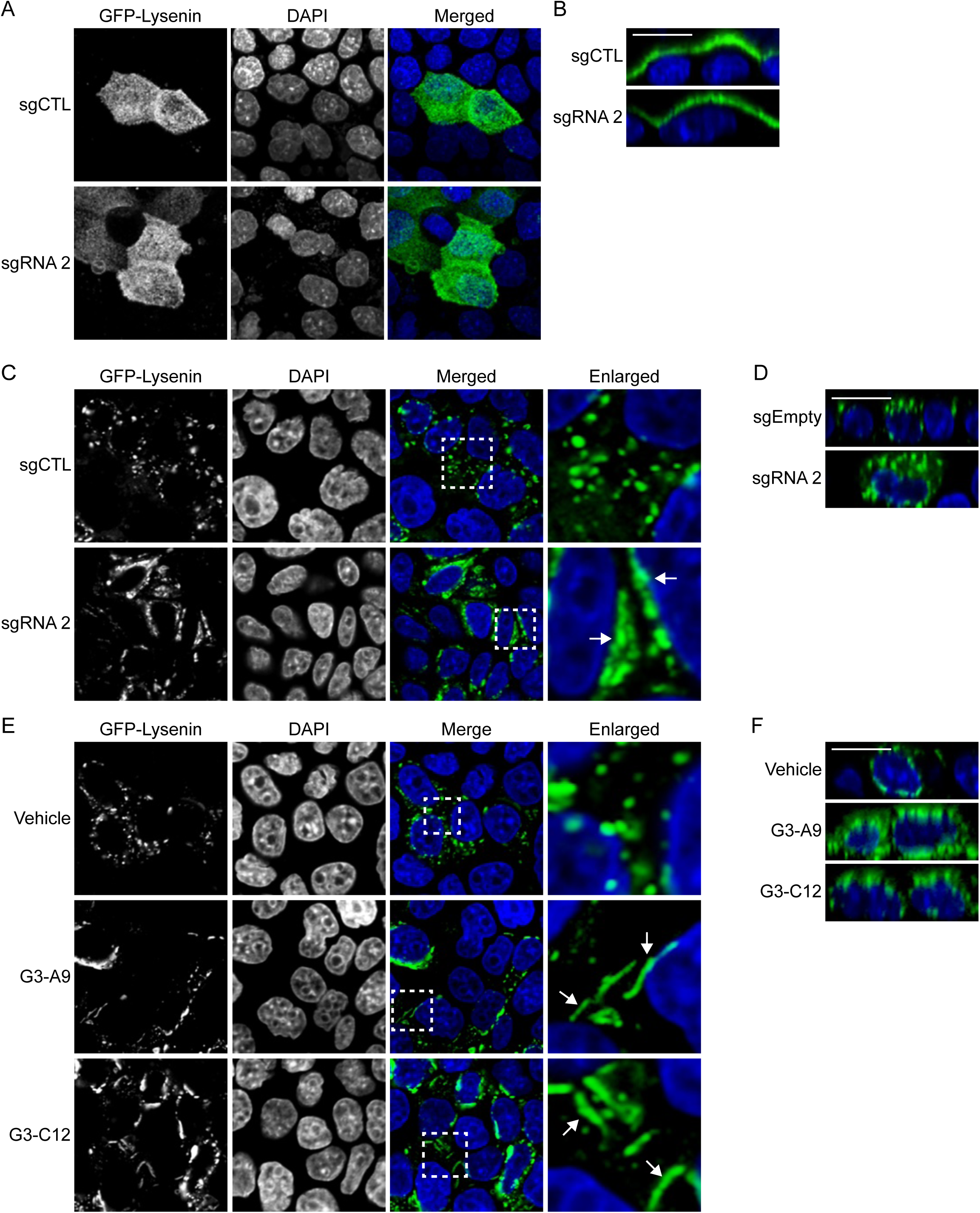
*Gal-3* KO and CRD inhibition perturb SM intracellular distribution. (A –. **D)** *Gal-3* KO MDCK cells that were un-permeabilized (A and B) or permeabilized (C and D) with saponin for 30 min were stained with 15 μg/ml GFP-lysenin for 15 min and DAPI for another 10 min at room temperature. (B) and (D) show the corresponding Z-stacks. **(E and F)** WT MDCK cells were treated with 50 μM Gal-3 CRD inhibitors for 48 h, permeabilized, and stained with GFP-lysenin and DAPI (E) with the corresponding Z-stack (F). The region outlined by the white square is shown at higher magnification. Arrowheads indicate the distribution of GFP-lysenin in elongated structures. Representative images from three independent experiments are shown. Scale bars = 10 μm.

### Gal-3 regulates the endocytic and recycling transport of SM, PtdSer and EGFR

We showed that *Gal-3* KO disrupts the intracellular distribution of PtdSer and SM. To further characterize the role of Gal-3 in the trafficking of these lipids, we examined the endocytic transport of SM and PtdSer by pulse-labeling cells with BODIPY-conjugated SM and PtdSer, TopFluor-SM and -PtdSer, respectively (Cho et al., 2012; Kay et al., 2012). Briefly, *Gal-3* KO or control MDCK cells were labeled with TopFluor-SM or - PtdSer for 10 min at 4 °C. Cells were washed and incubated at 37 °C, fixed at the indicated time points, and imaged by confocal microscopy. TopFluor-SM was predominantly localized at the PM after labeling (t = 0 min) and was internalized by 15 min in both control and *Gal-3* KO cell lines (Fig. 6A). While TopFluor-SM started to return to the PM at 45 min in control cells, it remained in vesicular structures at 45 min and started to return to the PM at 60 min in *Gal-3* KO cells. A similar distribution pattern was observed in both cell lines at 120 min. TopFluor-PtdSer was internalized from the PM to vesicular structures at 15 min and distributed extensively to cellular membranes after 45 min in control cells. In *Gal-3* KO cells, TopFluor-PtdSer remained at the PM at 15 min and was internalized at 30 min, followed by distribution to endomembranes after 60 min (Fig. 6B). Together, these data indicate that *Gal-3* KO delays the recycling of SM back to the PM without delaying its initial internalization, whereas for PtdSer, *Gal-3* KO delays both initial internalization and subsequent distribution to endomembranes.

**Figure 6.**
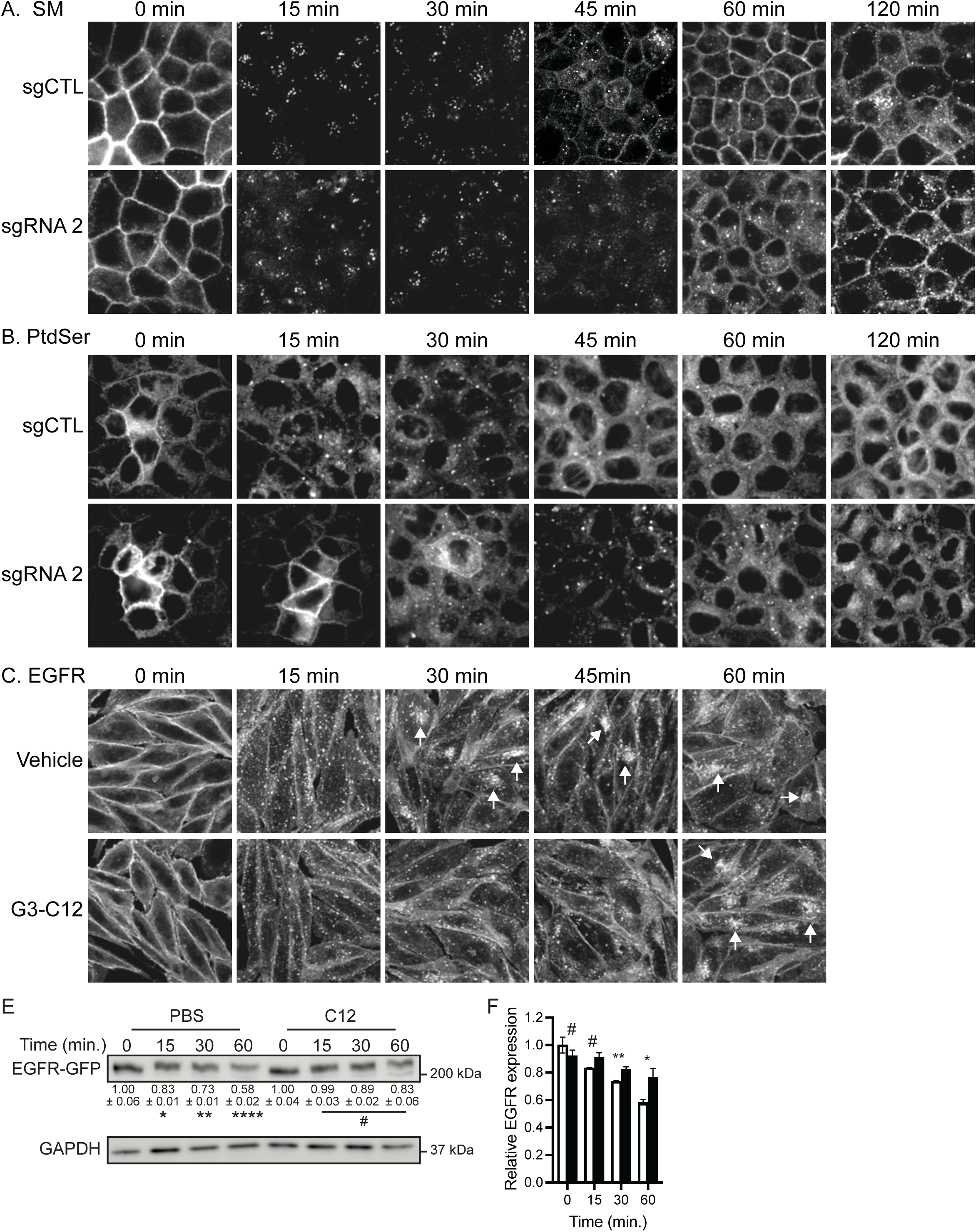
Gal-3 regulates the endocytic and recycling transport of SM, PtdSer and EGFR. (A and. **B)** *Gal-3* KO MDCK cells were pulse-labeled with 5 μM TopFluor-SM **(A)** or -PtdSer **(B)** in complete growth media for 10 min on ice at 4 °C. Cells were then washed with ice-cold PBS, refed with complete growth media and incubated at 37 °C for the indicated time. Cells were fixed and imaged by confocal microscopy. Representative images from three independent experiments are shown. Scale bars = 10 μm. **(C)** CHO-K1 cells stably overexpressing EGFR-GFP were treated with a Gal-3 CRD inhibitor for 48 h, and serum-starved for another 2 h in the presence of inhibitor. Cells were further incubated with 50 ng/ml human EGF for 15 min on ice, washed with ice-cold PBS and incubated with serum-free growth medium at 37 °C for the indicated time. Cells were fixed, stained with DAPI and imaged by confocal microscopy. Representative images are shown from three independent experiments. DAPI images are shown in Fig. S3. Scale bars = 10 μm. **(D)** Cell lysates from (C) were blotted with an anti-GFP antibody. Representative immunoblots are shown. **(E)** Quantification of (D). Values indicate the mean EGFR-GFP levels ± S.E.M. relative to vehicle-treated cells at t = 0, from three independent experiments. Significant differences between time points were assessed by one-way ANOVA, and between vehicle- and inhibitor-treated cells at each time point by Student’s t-tests (* p < 0.05, ** p < 0.01, # – not significant).

Next, we examined the role of Gal-3 in the endocytic transport of the epidermal growth factor receptor (EGFR). Chinese hamster ovary (CHO) cells, which have undetectable EGFR under basal conditions (Krug et al., 2002), were engineered to stably express EGFR-GFP and were treated with the Gal-3 CRD inhibitor for 48 h. Cells were then serum-starved, incubated with EGF for 15 min on ice, and EGFR-GFP trafficking was studied by time-lapse confocal microscopy. In control cells, EGFR-GFP was predominantly localized to the PM at 0 min followed by internalization at 15 min (Figs. 6C). EGFR-GFP started to accumulate in the perinuclear region, presumably recycling endosomes, at 30 min, consistent with a previous study (Baumdick et al., 2015), and was extensively distributed at 60 min. In Gal-3 CRD-inhibited cells, while EGFR-GFP was internalized at 15 min, its accumulation in the perinuclear region was observed only from 60 min after the stimulation, indicative of delayed post-endocytic trafficking of EGFR. Furthermore, our immunoblot data showed that Gal-3 CRD inhibition blocked EGF-induced EGFR degradation (Figs. 6D and E). Taken together, our data demonstrate that Gal-3 CRD promotes the endocytic and recycling trafficking of SM, PtdSer and EGFR, and that it is required for EGF-induced EGFR degradation.

## Discussion

In this study, we demonstrate that *Gal-3* KO or CRD inhibition depletes PtdSer and KRAS from the PM and disrupts their nanoscale spatial organization, and that exogenous PtdSer fully restores PM PtdSer while partially restoring KRAS (Fig. 3A). It has been reported that the Gal-3 CRD directly binds the farnesyl moiety of GTP-bound KRAS, and that blocking this interaction dissociates KRAS from the PM (Elad-Sfadia et al., 2004; Levy et al., 2010; Shalom-Feuerstein et al., 2008). Thus, our study identifies a second, lipid-mediated route of Gal-3–KRAS regulation, in which Gal-3 promotes KRAS PM binding through a direct CRD-mediated interaction and by maintaining PM PtdSer content. To identify the mechanism of Gal-3-mediated PM PtdSer maintenance, we examined lipids already known to control PM PtdSer content. PM PI4P, which drives ORP5/8-mediated PtdSer delivery at ER–PM contact sites (Chung et al., 2015; Moser von Filseck et al., 2015), PM cholesterol, whose transbilayer distribution is interdependent with that of PtdSer (Cho et al., 2016), and outer-leaflet GM3, proposed to support inner-leaflet PtdSer by cross-bilayer coupling (Liu et al., 2023; Raghupathy et al., 2015), were all unchanged (Figs. 3B – E). In addition, total PtdSer was unchanged while SM and Cer were reciprocally altered (Figs. 4A and B), and the endocytic and recycling transport of PtdSer, SM and EGFR was delayed (Figs. 5 and 6). Together, these data suggest that Gal-3 maintains PM PtdSer through vesicular transport.

In the extracellular space, the N-terminal domain (NTD) of monomeric Gal-3 engages in a transient intramolecular interaction with the CRD (Halimi et al., 2014; Ippel et al., 2016). When the CRD binds multivalent glycans of galactose-containing glycoproteins and glycolipids, it induces a conformational change that releases the NTD from the CRD, promoting NTD-mediated Gal-3 self-oligomerization (Halimi et al., 2014; Lepur et al., 2012; Zhao et al., 2021). Secreted Gal-3 re-enters cells via clathrin-independent endocytosis (CIE) and cycles through endosomal and recycling compartments (Straube et al., 2013). Within endosomes, Gal-3 oligomerization creates a dense Gal-3 lattice, which crosslinks glycosylated membrane components into microdomains. By scaffolding these luminal microdomains, Gal-3 structurally supports the cross-membrane clustering required for cytosolic adaptor protein binding to the endosome (Lakshminarayan et al., 2014; Nabi et al., 2015; Partridge et al., 2004). One example is ALG-2-interacting protein X (Alix), which recruits ESCRT-III to ubiquitinated endosomal cargo for delivery to multivesicular bodies and lysosomes (Christ et al., 2017; Raiborg and Stenmark, 2009). EGF stimulation promotes formation of Gal-3/Alix/EGFR complexes in CIE-derived endosomes, leading to lysosomal degradation, and silencing *Gal-3* or *Alix* delays EGFR internalization and subsequent endosomal sorting (Liu et al., 2012; Mercier et al., 2016; Wang et al., 2014). Consistent with this, we demonstrated that CRD inhibition delays the perinuclear accumulation of internalized EGFR-GFP and blocks EGF-induced EGFR degradation (Figs. 6C – E). When the Gal-3 CRD is blocked, endosomal Gal-3 cannot assemble the lattice. Alix, in turn, is not recruited to CIE-derived endosomes, and EGF-stimulated EGFR is retained in endosomes rather than undergoing lysosomal degradation.

In this context, we propose that Gal-3 maintains PM PtdSer and SM levels by regulating their intracellular trafficking. *Gal-3* KO delays the return of SM to the PM without perturbing its internalization, whereas both processes are delayed for PtdSer (Figs. 6A and B). While the exact endocytic mechanism for PtdSer remains to be fully elucidated (Kay and Fairn, 2019), endocytosed PtdSer returns to the PM via recycling endosomes (Cho et al., 2012; Uchida et al., 2011). In addition, excess SM has been proposed to impair PtdSer recycling out of the endolysosomal network, causing PtdSer to accumulate in endosomal structures and depleting PtdSer from the PM (Cho et al., 2012; Garrido et al., 2020; Maekawa et al., 2016). Thus, it is plausible that the SM elevation we observe, resulting from reduced neutral SMase activity and SMPD2 protein levels (Fig. 4), contributes directly to the impaired PtdSer recycling and the resulting loss of PM PtdSer. However, the delayed PtdSer internalization is not explained by this mechanism.

PM SM is internalized largely by caveolae (Nusshold et al., 2013), and in MDCK cells Gal-3 accumulates in caveolin-1- and flotillin-1-positive patches at the apical membrane before entering the cell by a cholesterol-dependent, non-clathrin route (Straube et al., 2013). Also, a carbohydrate-binding-deficient Gal-3 mutant (R186S) fails to associate with detergent-resistant membranes, indicating that CRD activity is required for recruitment of Gal-3 into these microdomains (Straube et al., 2013). Thus, CRD inhibition may disable a caveolae-associated recycling process, delaying the return of SM to the PM and driving its accumulation in perinuclear structures (Fig. 5). Alternatively, since PtdSer is required for caveola assembly (Hirama et al., 2017), the perturbed caveolae could arise secondary to PM PtdSer depletion, and the two defects may constitute a feed-forward loop.

In this study, we identify the Gal-3 CRD as a key regulator of PM lipid composition and KRAS PM localization. We propose that blocking the Gal-3 CRD prevents endosomal Gal-3 from assembling the lattice, which delays the endocytic and recycling trafficking of EGFR, SM and PtdSer. The elevated SM further impairs PtdSer recycling back to the PM. These concomitantly deplete PM PtdSer contents, dissociating KRAS from the PM. Thus, Gal-3 promotes KRAS PM binding through the direct CRD-mediated KRAS interaction and by maintaining PM PtdSer contents. Elucidating how Gal-3 regulates intracellular lipid trafficking will provide deeper understanding of the broader role of Gal-3 in human disease.

## Methods and Materials

### Plasmids and Reagents

Galectin-3 antibody was purchased from Santa Cruz (Cat# sc-32790). The following antibodies were purchased from Proteintech: β-actin (Cat# 66009-1-Ig), GFP-tag (Cat# 60002-1-Ig) and GAPDH (Cat# 10494-1-AP). The galectin-3 peptide inhibitors, G3-A9 and G3-C12, were synthesized by Bio-Synthesis (Lewisville, TX) (Zou et al., 2005). The amino acid sequence of G3-A9 is PQNSKIPGPTFLDPH and for G3-C12 is ANTPCGPYTHDCPVKR. Both Gal-3 peptides were dissolved in sterile 1x phosphate-buffered saline (PBS). TopFluor-sphingomyelin (810265P-250UG) and TopFluor-phosphatidylserine (810283) were from Avanti Polar Lipids.

### Cell lines

MDCK and CHO cells were grown in Dulbecco’s Modified Eagle Medium (DMEM) containing high glucose (Gibco, Cat# 10569-044) and F-12K medium (ATCC, 30-2004), respectively. They were supplemented with 2 mM L-glutamine (CA009-010, GenDepot) and 10% fetal bovine serum (FBS) (16000-044, Life Technologies). Cells were grown at 37°C in a 5% CO_2_ humidified chamber. Cell lines were tested periodically for mycoplasma using the MycoAlert Mycoplasma Detection Kit (Lonza, LT07-318). EGFR-GFP-CHO-1 cells were cultured with F-12K medium supplemented with 10% FBS and 2 mM L-Glut.

### Generation of Gal-3 CRISPR/Cas9 cell lines

Single guide RNAs were designed to target one of two different sites on canine Gal-3. For sgRNA2 and 3, it is 5’-ACCAAGCTGGATAATATCTG-3’ and 5’-CTTGCTCTAGATTTCAAGAG-3’. Single-stranded DNA oligonucleotides were designed and synthesized to be used as cloning primers for each specific Gal-3 sgRNA target site. For Gal-3 sgRNA 2, forward 5’ CACCGACCAAGCTGGATAATATCTG-3’, and reverse 5’ AAACCAGATATTATCCAGCTTGGTC-3’. For Gal-3 sgRNA 3, forward 5’ CACCGCTTGCTCTAGATTTCAAGAG-3’ and reverse 5’ AAACCTCTTGAAATCTAGAGCAAGC-3’. Each Gal-3 CRISPR sgRNA primer pair was 5’-phosphorylated using 10 µM of the relevant forward and reverse sgRNA primer, 5 U of T4 polynucleotide kinase (PNK), 1 µl of 10x T4 Ligation buffer which contains ATP (do not use 10x T4 PNK buffer) in a 10 µl total volume at 37°C for 30 min and then annealed at 95°C for 5 min. Approximately 10 pmol of the annealed-phosphorylated oligos were then simultaneously digested-ligated into 25 ng of the pSpCas9(BB)-2A-Puro (PX459) vector using 2.5 U BbsI, 200 U T4 ligase, 1 µl 10x T4 ligation buffer in a 10 µl total volume for 30 cycles of digestion at 37°C for 5 min and ligation at 23°C for 5 min. Approximately 5 µl of each new Gal-3 CRISPR/Cas9 plasmid was used to transform 50 µl of DH5α bacterial cells, which were plated onto LB-ampicillin plates overnight. Gal-3 CRISPR transformants were screened the next day using colony PCR assay, which consisted of 10 µM of both the forward U6 sequencing primer (5’-GATACAAGGCTGTTAGAGAGATAATT-3’) and the relevant Gal-3 reverse primer given above, 3% DMSO, 3 µl of the dissolved relevant colony and 10 µl AmfiSure PCR Master Mix (P0311, GenDepot) in a 20 µL total volume according to the manufacturer’s protocol. Products from the colony PCR Assay were electrophoresed on a 2% agarose gel and Gal-3 CRISPR/Cas9 clones that contained the correct amplicon were used to prepare plasmid DNA mini-preps from 5 ml bacterial cultures grown overnight in the presence of 100 µg/ml ampicillin. Gal-3 CRISPR/Cas9 plasmids were sequenced using the forward U6 sequencing primer.

MDCK cells were plated into 6-well plates on day 1. On day 2, cells were transfected with 4 µg of the relevant PX459 Gal-3 CRISPR sgRNA plasmid DNA and Lipofectamine 2000 (ThermoFisher), according to the manufacturer’s instructions. On day 3, transfection-containing reagents on cells were replaced with fresh complete DMEM growth media containing 2 µg/ml puromycin, which was reduced to 1 µg/ml puromycin on day 10. RNA extracts were prepared from polyclonal populations of cells that were grown for 2 days in 6-well plates which were then reverse transcribed into cDNA and used for PCR-based screening for corroboration of CRISPR/Cas9 indel events surrounding the targeted sgRNA sites. At least 10 monoclonal cell lines were generated for either sgRNA2 or sgRNA3 using limiting dilution in order to generate subclones with well-defined CRISPR/Cas9-mediated indels from a relatively homogenous population of cells. Sanger sequencing of PCR products was performed for both polyclonal and monoclonal cell lines to confirm the presence of CRISPR/Cas9-mediated indels. For both sgRNA2 and sgRNA3, clones carrying a single-nucleotide insertion that caused a frameshift and introduced a premature termination codon within the Gal-3 CRD were selected for use in this study. Also, MDCK cells were transfected with empty PX459 vector to be used as a negative control.

### RNA isolation and RT-PCR

RNA was extracted from cells grown in a single 35 mm well using the RNeasy kit (74104, Qiagen) and according to the manufacturer’s instructions. Two micrograms of total RNA from each sample was used for the synthesis of first-strand cDNA using Superscript II reverse transcriptase (18064-014, ThermoFisher). Reverse transcription-PCR (RT-PCR) was performed in a 20 µl reaction containing 100 ng cDNA, 500 nM each of the relevant forward and reverse primers, 200 µM dNTPs, 2 µl of either the 5x Phusion GC Buffer or the 5x HF Buffer (depending on the percentage of GC content of the amplicon), 3-9% DMSO and 0.2 µl Phusion High-Fidelity DNA Polymerase (F-553L, ThermoFisher) according to this manufacturer’s recommendations. PCR products were separated on a 2% agarose gel containing Midori Green Advance Safe DNA Stain (MG04, Bulldog-Bio, Portsmouth, NH) in 1x TAE buffer and were visualized in the agarose gels using a blue LED illuminator. PCR products were excised from the gels and purified using the Genejet Gel Extraction Kit (K0692, ThermoFisher) according to the manufacturer’s protocol. The forward and reverse primers used to screen for the presence of the Gal-3 sg2 or the sg3 target sites are given above. Agarose gel- and column-purified PCR products were sent to Genewiz (South Plainfield, NJ) for Sanger sequencing analyses.

### Confocal Microscopy

MDCK cells were seeded on coverslips at 2.5 x 10^5^ on a 12-well plate. 72 h after the seeding, cells were washed twice with ice-cold 1x phosphate-buffered saline (PBS) and then fixed with 4% paraformaldehyde (Electron Microscopy Services, Cat #15710) followed by 50 mM NH_4_Cl. Slides were imaged using the Olympus FV1000 confocal microscope using a 60x objective. Images were quantified for co-localization using Manders’ coefficient on ImageJ software (version 1.52a).

### Electron microscopy (EM) and spatial mapping

Plasma membrane sheets were prepared and fixed as previously described (Cho et al., 2012; Garrido et al., 2020; Hancock and Prior, 2005; Prior et al., 2003). For univariate analysis, plasma membrane sheets were labeled with anti-green fluorescent protein (anti-GFP) antibody conjugated to 4.5-nm gold particles. Digital images of the immunogold-labeled plasma membrane sheets were taken in a transmission electron microscope. Intact 1 μm^2^ areas of the plasma membrane sheet were identified using ImageJ software, and the (*x*, *y*) coordinates of the gold particles were determined (Hancock and Prior, 2005; Prior et al., 2003). K-functions (Ripley, 1977) were calculated and standardized on the 99% confidence interval (CI) for univariate functions (Diggle, 2000; Hancock and Prior, 2005; Prior et al., 2003). In the case of univariate functions, a value of *L*(*r*) - *r* greater than the CI indicates significant clustering, and the maximum value of the function (*L*_max_) estimates the extent of clustering. Differences between replicated point patterns were analyzed by constructing bootstrap tests as described previously (Diggle, 2000; Plowman et al., 2005), and the statistical significance against the results obtained with 1,000 bootstrap samples was evaluated.

### Manders’ Coefficient

MDCK cells stably expressing two different fluorophores were imaged as stated above using a confocal microscope. Tagged image file format (.tiff) confocal images were converted to 8-bit files, and a threshold to a control pixel of each image was set. The fraction of mCherry-CAAX endomembrane marker that co-localized with either GFP-KRASG12V or GFP-LactC2 was calculated using the Manders Coefficient plugin downloaded from imagej.nih.gov.

### Lysenin staining

Lysenin staining was performed as preiously described (van der Hoeven et al., 2018). Briefly, cells used were on glass coverslips that were first fixed with 4% PFA and then permeabilized with 0.05% saponin in 1x PBS for 30 min at room temp. Permeable cells were then incubated with 15 µg/ml MBP-GFP-lysenin for 15 min at room temp and washed twice with 1x PBS. Cells were then incubated with 50 mM NH4Cl for 10 min at room temp, washed again twice with 1x PBS followed by an incubation with 1:2000 dilution of DAPI in PBS for 10 min in the dark. Cells were washed once with 1x PBS, once with water and then mounted onto glass microscope slides using Vectashield mounting media. Cells were imaged using a confocal microscope.

### Top-Fluor-SM and -PtdSer endocytosis

On day 1, 2.75x10^5^ MDCK WT CRISPR sgCTL or GAL-3 sgRNA2 cells were seeded onto glass coverslips in 12-well plates containing 1 ml complete growth media (DMEM, 10% FBS, 2 mM L-glutamine). On day 2, cells were washed twice with 1 ml sterile 1x PBS and then refed with 2 ml complete media. On day 3, TopFluor-SM or -PtdSer were initially dissolved in chloroform at 1 mg/ml and were dried overnight under a vacuum in an amber glass vial to remove the solvent. On day 4, the dried lipid of interest was then redissolved in 2 ml serum-free growth media (DMEM, 2 mM L-glutamine) using sonication for 1 h and then diluted to a final concentration of 5 µM using complete growth media, which was then kept on ice. At this point, cells in 12-well plates were placed on ice and gently washed twice with 1 ml of sterile, ice-cold 1x PBS in a refrigerated cold-room at 4°C. Cells were incubated for 10 min in 1 ml of 1x PBS and then the PBS was removed and replaced with 500 µl of 5 µM TopFluor-SM, or -PtdSer. Lipid-treated cells were incubated for 10 min on ice in the refrigerated cold-room. Next, cells were gently washed three times with 1 ml ice-cold 1x PBS in the cold room. The 0 min time point cells were fixed using 1 ml of 4% PFA, while cells for all other time points were fed with 2 ml of pre-warmed (37°C) complete growth media to initiate lipid endocytosis and immediately were placed back in the 37°C cell culture incubator for up to 120 min. After completion of each subsequent time point, cells were washed twice with 1 ml ice-cold 1x PBS and fixed using 1 ml of 4% PFA for 1 h. Cells were washed twice with 1 ml 1x PBS and then incubated with 50 mM NH_4_Cl for 10 min at room temp. Next, cells were again washed with PBS and then incubated with 1:2000 dilution of DAPI in PBS for 10 min at room temp to stain the nuclei. Cells were imaged using a confocal microscope.

### Preparing phosphatidylserine

1,2-Diacyl-sn-glycero-3-phospho-L-serine (P6641, Sigma) supplied in chloroform:methanol was dried under vacuum in a glass vial to remove the solvent, redissolved in MDCK growth medium by sonication, and diluted to give a final concentration of 10 μM.

### Western Blotting

Cells were washed twice with ice-cold 1x PBS. Cells were harvested in a lysis buffer consisting of 50mM Tris pH 7.5, 25 mM NaF, 5 mM MgCl_2_, 75 mM NaCl, 5 mM EGTA, 1 mM DTT, 100 μM NaVO_4_ and 1% NP-40. Protein concentrations were determined using the BCA protein assay. SDS-page and immunoblotting were performed using 20-25 μg of lysate from each sample group. Blots were detected using chemiluminescence on the Amersham Imager 600. Blots were quantified using ImageJ software.

### Cy5-annexin V-binding analyses

MDCK WT sgCTL or CRISPR/Cas9-Gal-3 sgRNA2 cells were incubated with Cy5-conjugated annexin V (559934, BD Pharmigen) according to the manufacturer’s protocol. A portion of each cell sample was incubated with 1 µM staurosporine for 6 h to induce apoptosis and flip PtdSer to the outer PM leaflet. Cy5-annexin V-positive cells were counted using a BD AccuriC6 analyzer with a 670/30 nm filter.

### Lipidomics

Cells were washed twice with 1 ml sterile 1x PBS and then 500 µl 0.25% trypsin-EDTA was applied to cells. Cells were trypsinized until all cells were loosened and then combined with ∼1.5 ml 1x Dulbecco’s phosphate-buffered saline (D-PBS). Cell concentrations were determined using trypan blue exclusion and a glass hemocytometer. For each sample, 1 x 10^6^ cells in 333 µl D-PBS was analyzed for lipid composition. Lipid extraction and analyses were conducted by Lipotype GmbH (Dresden, Germany) via electionspray ionization and tandem mass spectrometry, as previously described (Gerl et al., 2012; Sampaio et al., 2011).

### Lysosomal pH measurement

Either MDCK WT sgCTL and CRISPR/Cas9-Gal-3 sgRNA2 cells or MDCK WT cells were seeded in triplicate wells of a 96-well plate at a density of 1.56 x 10^4^ cells per well on Day 1. On Day 2, MDCK WT only cells were treated with vehicle or 50 µM Gal-3 G3-C12 peptide. At this same time, 1 µg/µl Lysosensor Yellow/Blue Dextran reagent was applied to the complete DMEM growth media that was covering the wells of MDCK WT sgCTL and CRISPR/Cas9-Gal-3 sgRNA2 cells and also MDCK WT cells treated with vehicle or 50 µM Gal-3 G3-C12 peptide. Forty-eight h later on Day 4, all wells of cells in the 96-well plate were carefully washed twice with 200 µl PBS and then 100 µl of PBS was added to the wells. The 96-well plate was read in a BioTek Synergy H1 microplate reader with excitation/emission at 355 nm/452 and 381 nm/521 nm. The mean ratio of emission 452 nm/521 nm was then calculated ± SEM for each sample (Garrido et al., 2020).

### Sphingomyelinase activity assay

The SMase activity assay was performed with Amplex Red SMase Assay Kit (A12220, Invitrogen) as described previously (Tan et al., 2019). Briefly for the neutral SMase assay, 10 μg of whole-cell lysates (at 1 μg/μl concentration) prepared in lysis buffer B without DTT was mixed with 40 μl of 1x reaction buffer (0.5 M Tris-HCl pH 7.4, 50 mM MgCl2), and loaded on a well of black 96-well plate. 50 μl of Amplex Red reaction mixture (100 μM Amplex Red reagent, 2 U/ml horseradish peroxidase, 0.2 U/ml choline oxidase, 8 U/ml alkaline phosphatase, and 0.5 mM sphingomyelin in 1x reaction buffer) was added to each well, and the plate was incubated in dark at 37 °C for 60 min. The fluorescence was measured using BioTek Synergy H1 microplate reader (excitation λ = 540 nm, emission λ = 590 nm). 0.1 U/ml SMase and 1x reaction buffer were used as a positive and a negative control, respectively. For the acid SMase assay, 10 μg of whole-cell lysates (at 1 μg/μl concentration) prepared in lysis buffer B without DTT was mixed with 40 μl of low-pH buffer (50 mM sodium acetate, pH 5.0), and loaded on a well of black 96-well plate. 5 μl of 5 mM sphingomyelin was added to each well, and the plate was incubated in dark in 37 °C for 1 h. After the incubation, the pH was raised to 7.4 by adding 50 μl of Amplex Red reaction mixture [100 μM Amplex Red reagent, 2 U/ml horseradish peroxidase, 0.2 U/ml choline oxidase, 8 U/ml alkaline phosphatase in high-pH buffer (100 mM Tris-HCl, pH 8.0)]. The plate was further incubated in the dark in 37 °C for 60 min, and the fluorescence was measured using BioTek Synergy H1 microplate reader (excitation λ = 540 nm, emission λ = 590 nm). 0.1 U/ml SMase and low-pH buffer were used as a positive and negative control, respectively.

### Statistics

GraphPad Prism software (v.8.4.0) was used for one-way ANOVA and Student’s *t*-test. Bootstrap tests for the spatial mapping data were performed as described above. Lipidomics p values were corrected for multiple comparison across lipid species using the Benjamini-Hochberg procedure.

## Supporting information

Combined Supp Fig

## Acknowledgements

We would like to thank Karen Henkels for her contribution to this work. This work was supported by NIH grants (R15 GM163279) to KJC, and (R01 GM151280) and a CPRIT grant (RP250213) to JL and JFH.

