## Supplementary material for "The galectin-3 carbohydrate recognition domain maintains plasma membrane phosphatidylserine required for KRAS function": Combined Supp Fig

#### Supplementary Figure Legends

**Figure S1. *Gal-3* KO depletes KRAS and PtdSer from the PM. (A)** *Gal-3* KO MDCK cells co-expressing mCherry-CAAX and GFP-LactC2 were seeded on glass coverslips, fixed with 4% PFA and imaged by confocal microscopy. Values represent mean ± S.E.M. (n = 3) of the fraction of mCherry fluorophore co-localized with GFP fluorophore, calculated by Manders' coefficient. Representative images from three independent experiments are shown. **(B)** Intact basal PM sheets prepared from *Gal-3* KO MDCK cells stably expressing GFP-KRASG12V were labeled with 4.5 nm gold-conjugated anti-GFP antibodies and visualized by EM. The graph shows a mean number of gold particles ± S.E.M. (n ≥ 15). Representative images are shown.

**Supplement Figure 2. *Gal-3* KO and CRD inhibition disrupt cellular lipid levels.** MDCK cells stably expressing GFP-KRASG12V that are *Gal-3* KO or treated with G3-C12 for 48 h were prepared and whole cell lipid levels were measured from three independent experiments. A heat map was constructed to quantify the changes of different lipid species in *Gal-3* KO or CRD inhibited cells. Each row represents a different species of lipid, while each column represents a single sample. The scaled expression values of each lipid measured are plotted in red-blue color $\log_2$ scale. In relation to control cells, red and blue colored tiles indicate higher and lower lipid contents, respectively. **(A)** CE: cholesteryl ester, **(B)** HexCer: hexosylceramide, **(C)** PA: phosphatidic acid, **(D)** LPC: lysophosphatidylcholine, LPC O-: ether-linked LPC **(E)** LPE: lysophosphatidylethanolamine, LPE O-: ether-linked LPE **(F)** DAG: diacylglycerol, **(G)** CL: cardiolipin, **(H)** PI: phosphatidylinositol, **(I)** PG: phosphatidylglycerol, **(J)** PC – phosphatidylcholine, PC O-: ether-linked PC, and **(K)** PE – phosphatidylethanolamine, PE O- – ether-linked PE.

**Supplement Figure 3. Gal-3 CRD inhibition delays the post-endocytic trafficking of EGFR.** CHO-K1 cells stably overexpressing EGFR-GFP were treated with vehicle (PBS) (A) or 50 μM G3-C12 (B) for 48 h, and serum-starved for another 2 h in the continued presence of vehicle or inhibitor. Cells were further incubated with 50 ng/ml human EGF for 15 min on ice, washed with ice-cold PBS and incubated with serum-free growth medium at 37 °C for the indicated time. Cells were fixed, stained with DAPI and imaged by confocal microscopy. Representative images are shown from three independent experiments.

Figure S1

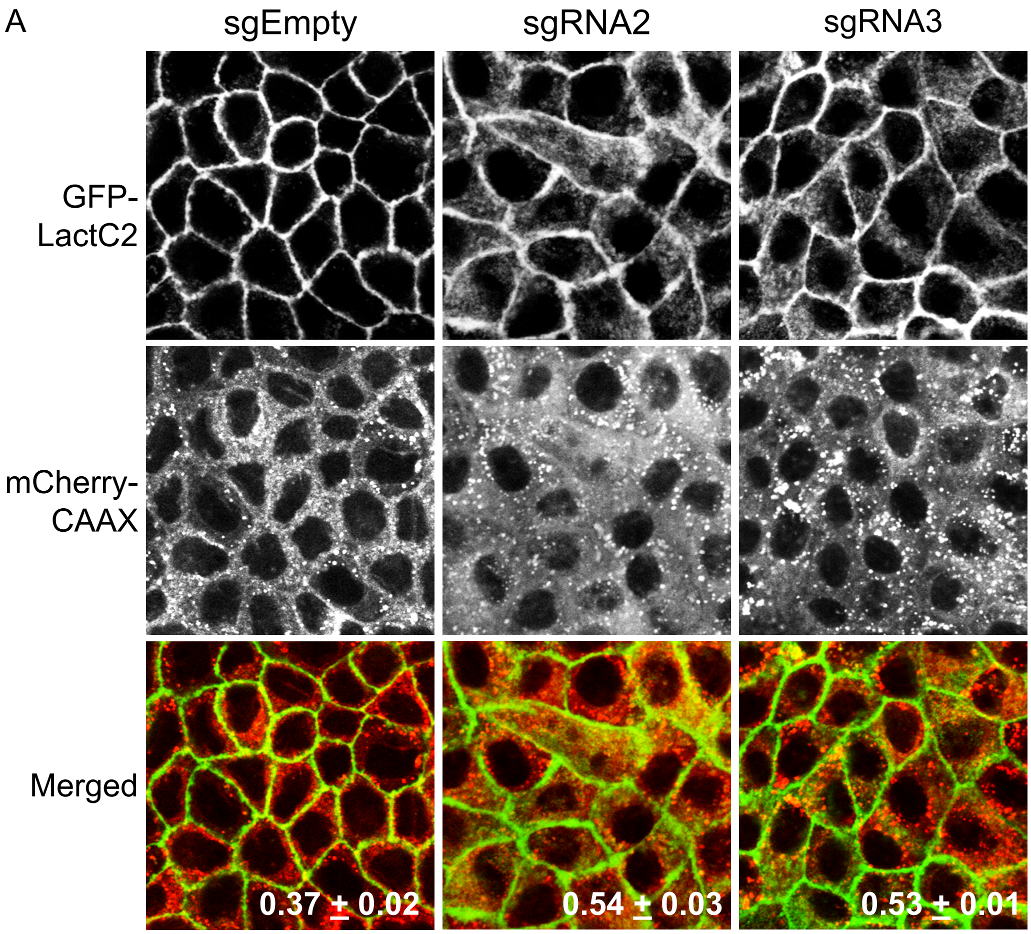
A
sgEmpty
sgRNA2
sgRNA3
GFP-LactC2
mCherry-CAAX
Merged
0.37 ± 0.02
0.54 ± 0.03
0.53 ± 0.01

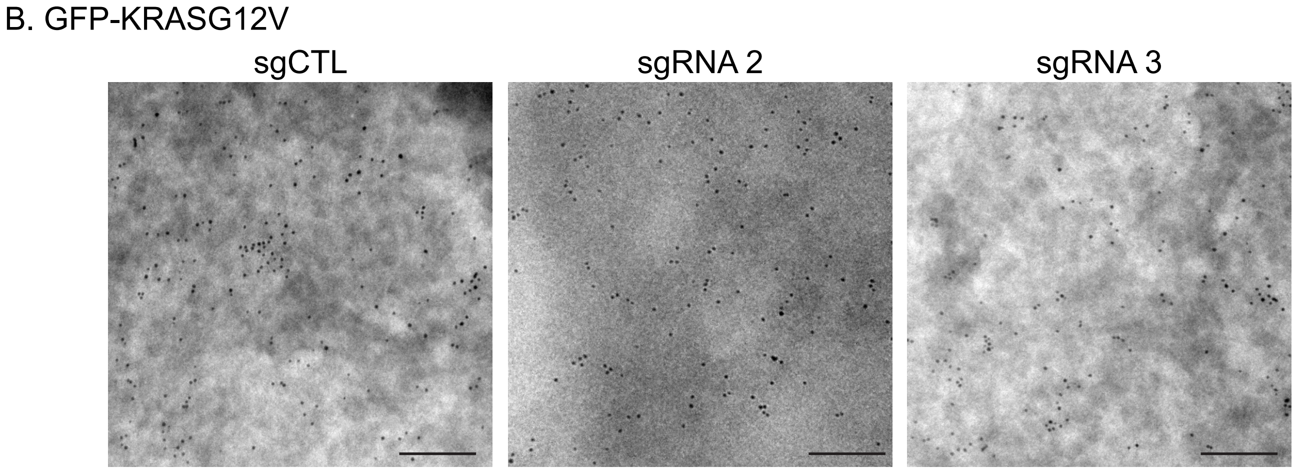
B. GFP-KRASG12V
sgCTL
sgRNA 2
sgRNA 3

Fig S2

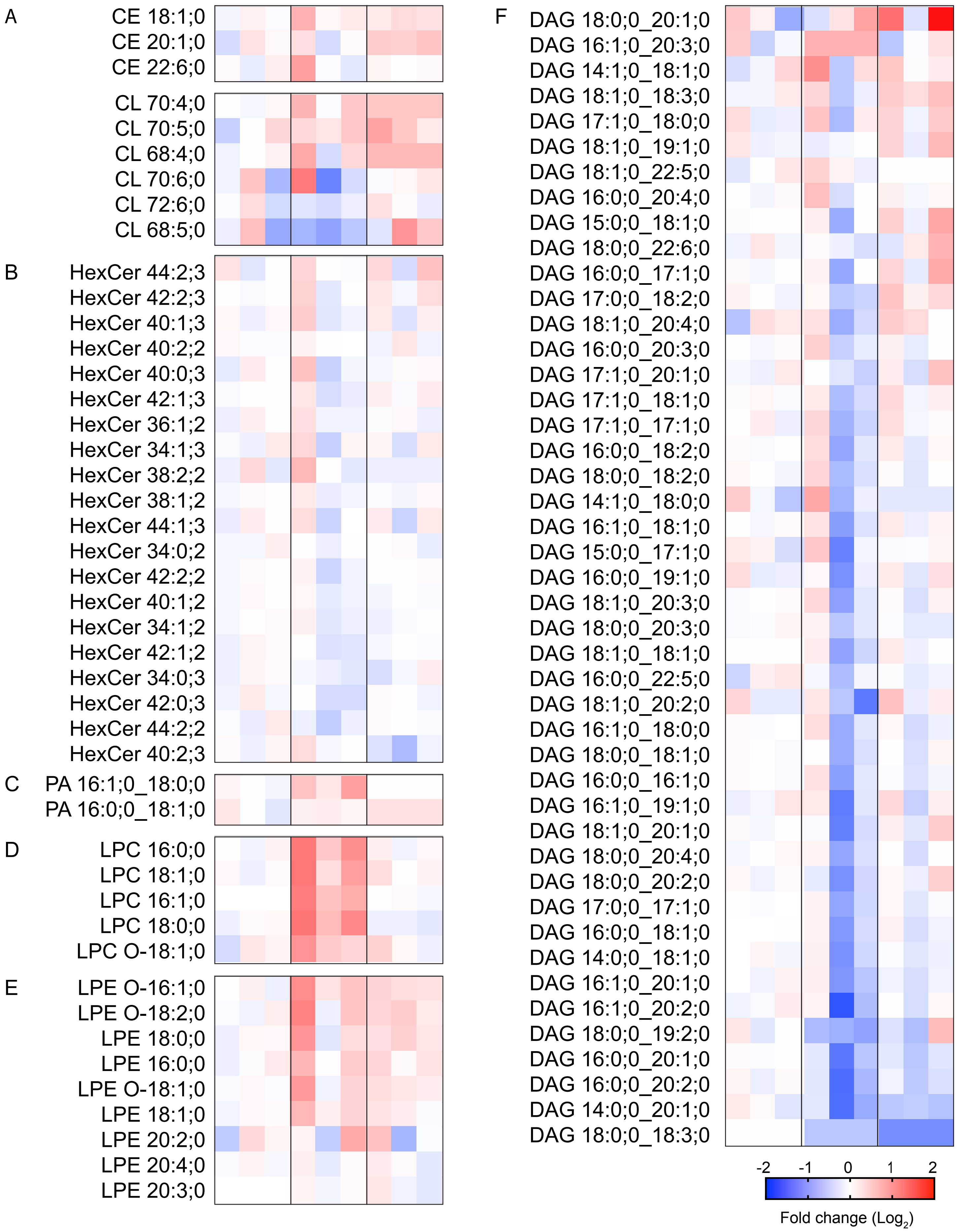
A
CE 18:1;0
CE 20:1;0
CE 22:6;0
CL 70:4;0
CL 70:5;0
CL 68:4;0
CL 70:6;0
CL 72:6;0
CL 68:5;0
B
HexCer 44:2;3
HexCer 42:2;3
HexCer 40:1;3
HexCer 40:2;2
HexCer 40:0;3
HexCer 42:1;3
HexCer 36:1;2
HexCer 34:1;3
HexCer 38:2;2
HexCer 38:1;2
HexCer 44:1;3
HexCer 34:0;2
HexCer 42:2;2
HexCer 40:1;2
HexCer 34:1;2
HexCer 42:1;2
HexCer 34:0;3
HexCer 42:0;3
HexCer 44:2;2
HexCer 40:2;3
C
PA 16:1;0_18:0;0
PA 16:0;0_18:1;0
D
LPC 16:0;0
LPC 18:1;0
LPC 16:1;0
LPC 18:0;0
LPC O-18:1;0
E
LPE O-16:1;0
LPE O-18:2;0
LPE 18:0;0
LPE 16:0;0
LPE O-18:1;0
LPE 18:1;0
LPE 20:2;0
LPE 20:4;0
LPE 20:3;0
F
DAG 18:0;0_20:1;0
DAG 16:1;0_20:3;0
DAG 14:1;0_18:1;0
DAG 18:1;0_18:3;0
DAG 17:1;0_18:0;0
DAG 18:1;0_19:1;0
DAG 18:1;0_22:5;0
DAG 16:0;0_20:4;0
DAG 15:0;0_18:1;0
DAG 18:0;0_22:6;0
DAG 16:0;0_17:1;0
DAG 17:0;0_18:2;0
DAG 18:1;0_20:4;0
DAG 16:0;0_20:3;0
DAG 17:1;0_20:1;0
DAG 17:1;0_18:1;0
DAG 17:1;0_17:1;0
DAG 16:0;0_18:2;0
DAG 18:0;0_18:2;0
DAG 14:1;0_18:0;0
DAG 16:1;0_18:1;0
DAG 15:0;0_17:1;0
DAG 16:0;0_19:1;0
DAG 18:1;0_20:3;0
DAG 18:0;0_20:3;0
DAG 18:1;0_18:1;0
DAG 16:0;0_22:5;0
DAG 18:1;0_20:2;0
DAG 16:1;0_18:0;0
DAG 18:0;0_18:1;0
DAG 16:0;0_16:1;0
DAG 16:1;0_19:1;0
DAG 18:1;0_20:1;0
DAG 18:0;0_20:4;0
DAG 18:0;0_20:2;0
DAG 17:0;0_17:1;0
DAG 16:0;0_18:1;0
DAG 14:0;0_18:1;0
DAG 16:1;0_20:1;0
DAG 16:1;0_20:2;0
DAG 18:0;0_19:2;0
DAG 16:0;0_20:1;0
DAG 16:0;0_20:2;0
DAG 14:0;0_20:1;0
DAG 18:0;0_18:3;0
-2
-1
0
1
2
Fold change (Log2)

Fig S2 Cont'd

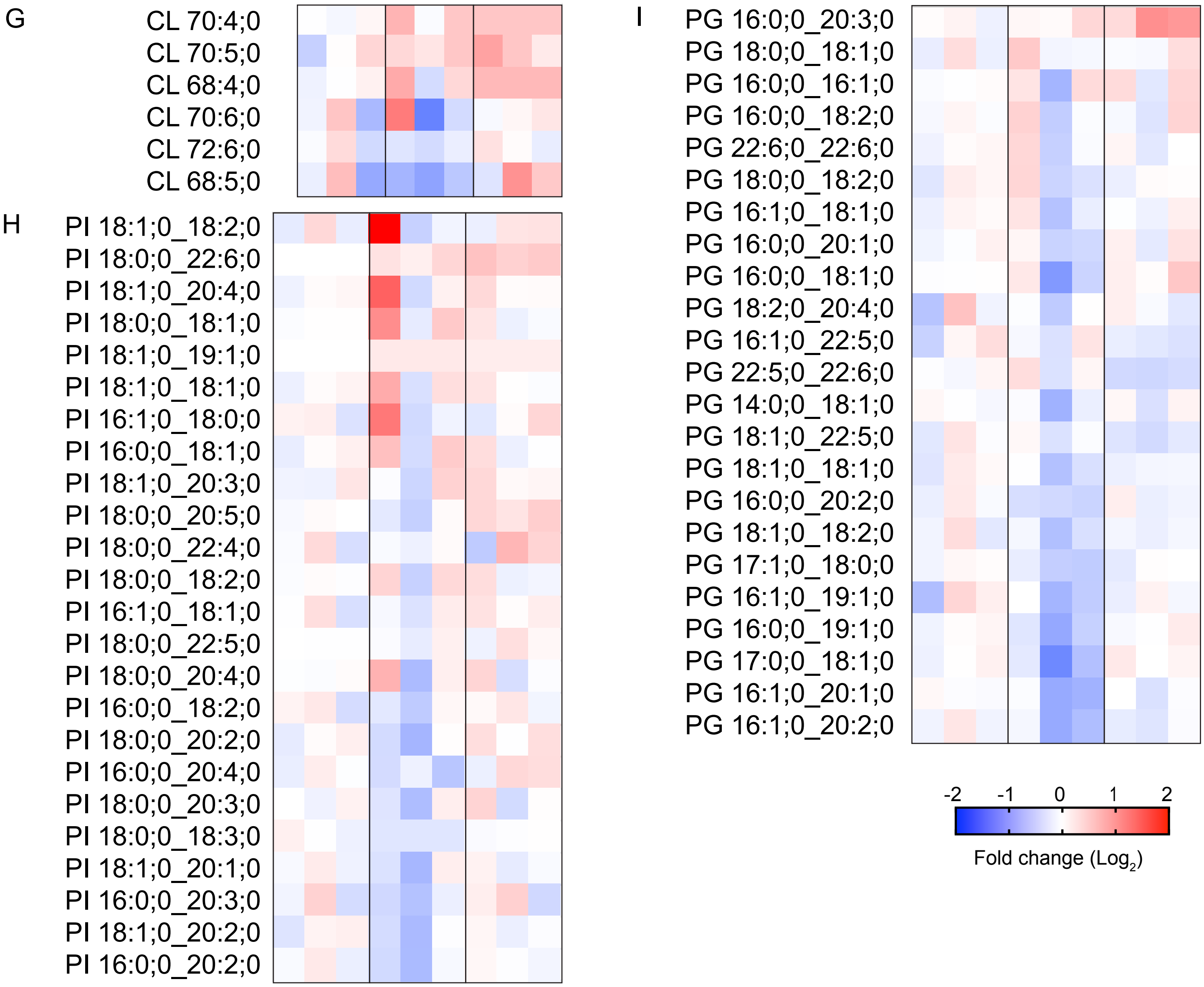
G
CL 70:4;0
CL 70:5;0
CL 68:4;0
CL 70:6;0
CL 72:6;0
CL 68:5;0
H
PI 18:1;0_18:2;0
PI 18:0;0_22:6;0
PI 18:1;0_20:4;0
PI 18:0;0_18:1;0
PI 18:1;0_19:1;0
PI 18:1;0_18:1;0
PI 16:1;0_18:0;0
PI 16:0;0_18:1;0
PI 18:1;0_20:3;0
PI 18:0;0_20:5;0
PI 18:0;0_22:4;0
PI 18:0;0_18:2;0
PI 16:1;0_18:1;0
PI 18:0;0_22:5;0
PI 18:0;0_20:4;0
PI 16:0;0_18:2;0
PI 18:0;0_20:2;0
PI 16:0;0_20:4;0
PI 18:0;0_20:3;0
PI 18:0;0_18:3;0
PI 18:1;0_20:1;0
PI 16:0;0_20:3;0
PI 18:1;0_20:2;0
PI 16:0;0_20:2;0
I
PG 16:0;0_20:3;0
PG 18:0;0_18:1;0
PG 16:0;0_16:1;0
PG 16:0;0_18:2;0
PG 22:6;0_22:6;0
PG 18:0;0_18:2;0
PG 16:1;0_18:1;0
PG 16:0;0_20:1;0
PG 16:0;0_18:1;0
PG 18:2;0_20:4;0
PG 16:1;0_22:5;0
PG 22:5;0_22:6;0
PG 14:0;0_18:1;0
PG 18:1;0_22:5;0
PG 18:1;0_18:1;0
PG 16:0;0_20:2;0
PG 18:1;0_18:2;0
PG 17:1;0_18:0;0
PG 16:1;0_19:1;0
PG 16:0;0_19:1;0
PG 17:0;0_18:1;0
PG 16:1;0_20:1;0
PG 16:1;0_20:2;0
-2
-1
0
1
2
Fold change (Log2)

### Figure S2. Cont'd

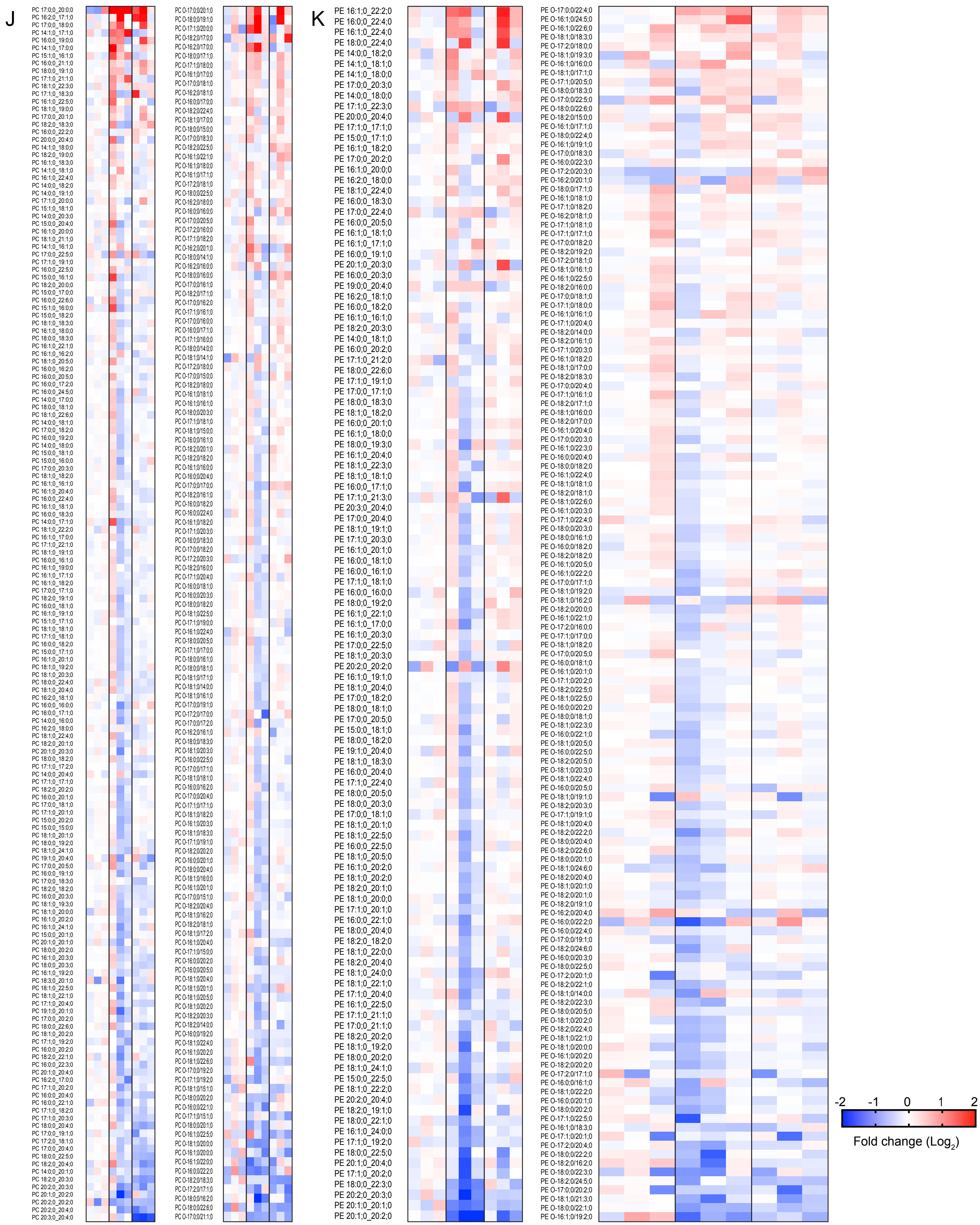

Figure S3

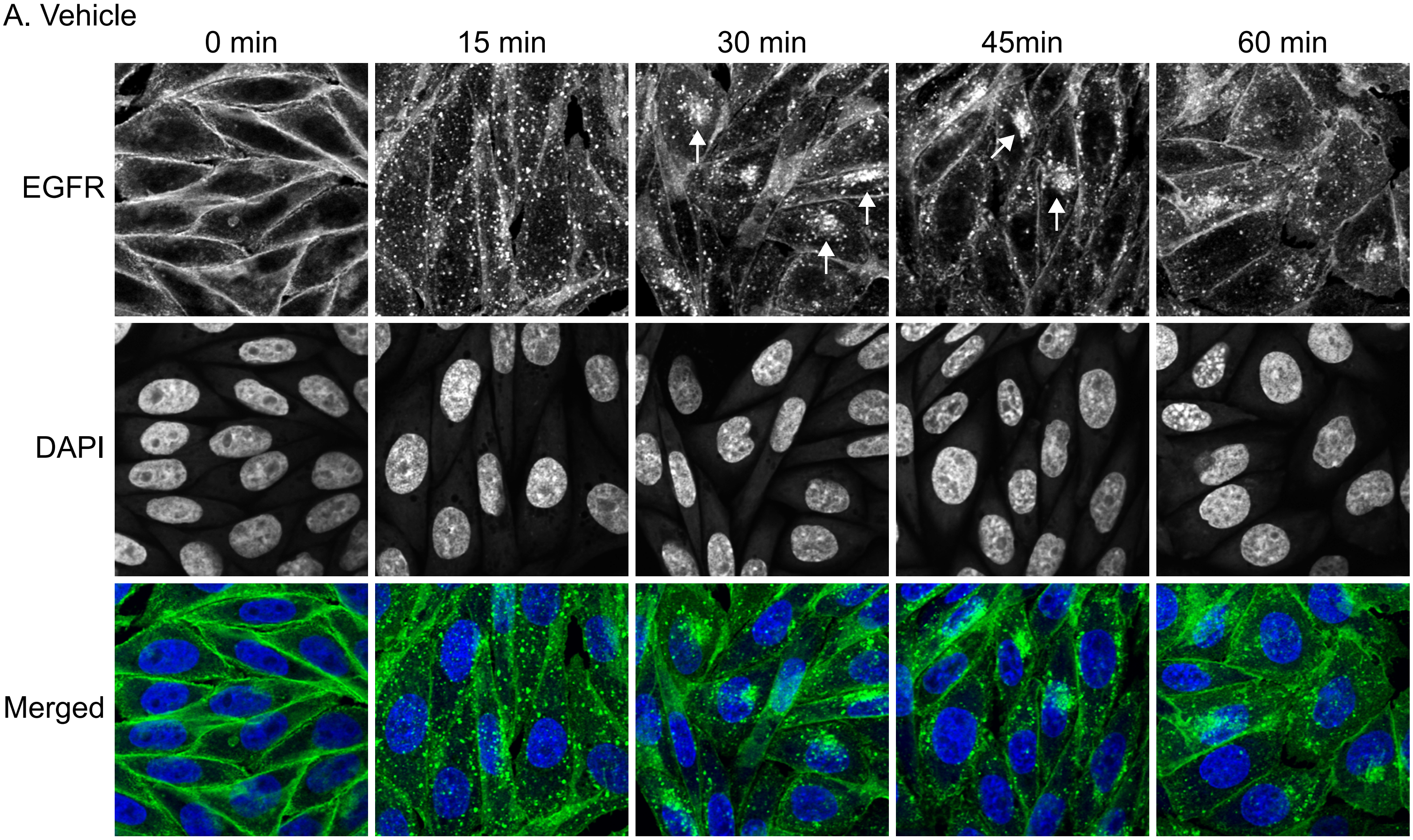
A. Vehicle
0 min
15 min
30 min
45min
60 min
EGFR
DAPI
Merged

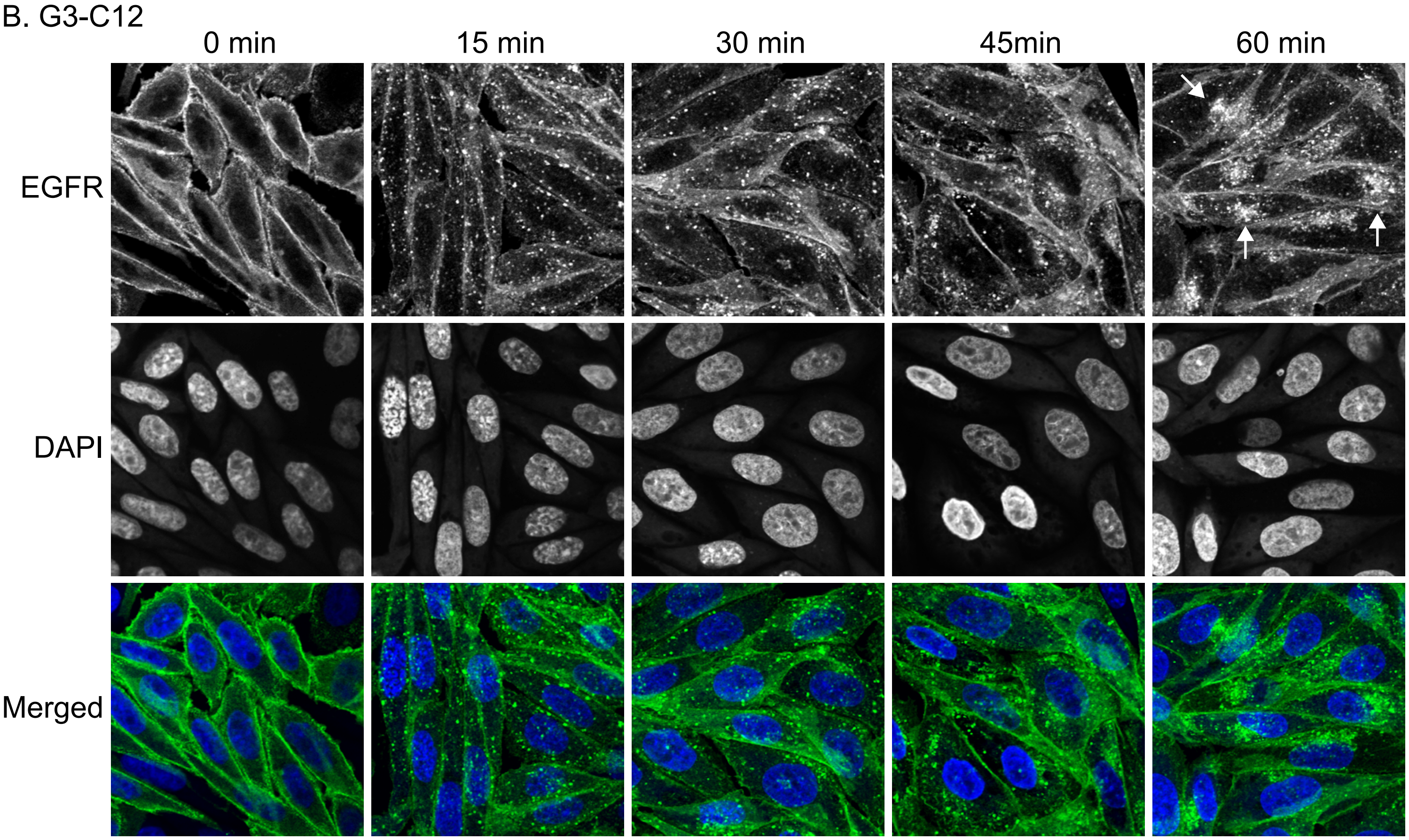
B. G3-C12
0 min
15 min
30 min
45min
60 min
EGFR
DAPI
Merged
